# Deficiency in the endocytic adaptor protein PHETA1/2 impairs renal and craniofacial development

**DOI:** 10.1101/727578

**Authors:** Kristin M. Ates, Tong Wang, Trevor Moreland, Rajalakshmi Veeranan-Karmegam, Priya Anand, Wolfgang Wenzel, Hyung-Goo Kim, Lynne A. Wolfe, Joshi Stephen, David R. Adams, Thomas Markello, Cynthia J. Tifft, William A. Gahl, Graydon B. Gonsalvez, May Christine Malicdan, Heather Flanagan-Steet, Y. Albert Pan

**Author notes:** Correspondence should be addressed: Dr. Y. Albert Pan,. **Conflict of Interest:** No competing interests declared.

## Abstract

A critical barrier in the treatment of endosomal and lysosomal diseases is the lack of understanding of the *in vivo* functions of the putative causative genes. We addressed this by investigating a key pair of endocytic adaptor proteins, PH domain containing endocytic trafficking adaptor 1 and 2 (PHETA1/2, also known as FAM109A/B, Ses1/2, IPIP27A/B), which interact with the protein product of *OCRL*, the causative gene for Lowe syndrome. Here we conducted the first study of PHETA1/2 *in vivo*, utilizing the zebrafish system. We found that impairment of both zebrafish orthologs, *pheta1* and *pheta2*, disrupted endocytosis and ciliogenesis. In addition, *pheta1/2* mutant animals exhibited reduced jaw size and delayed chondrocyte maturation, indicating a role in craniofacial development. Deficiency of *pheta1/2* resulted in dysregulation of cathepsin K, which led to an increased abundance of type II collagen in craniofacial cartilages. The abnormal renal and craniofacial phenotypes in the *pheta1/2* mutant animals were consistent with the clinical presentations of a patient with a *de novo* arginine (R) to cysteine (C) variant (R6C) of PHETA1. Expressing the patient-specific variant in zebrafish exacerbated craniofacial deficits, suggesting that the R6C allele acts in a dominant-negative manner. Together, these results provide insights into the *in vivo* roles of PHETA1/2 and suggest that the R6C variant is contributory to the pathogenesis of disease in the patient.

## Introduction

Endocytic trafficking is essential for a variety of biological processes, including nutrient uptake, cell signaling, and cellular morphogenesis (Doherty and McMahon, 2009). This diversity in cellular functions is reflected in the broad range of pathologies associated with deficiencies in endocytic factors. For example, mutations in endocytic factors *dynamin 2* (*DNM2*) and *RAB7* result in Charcot-Marie-Tooth disease, a clinically and genetically heterogeneous group of peripheral neuropathies (Verhoeven et al., 2003; Züchner et al., 2005). Disruptions in endocytosis have been identified in autosomal recessive hypercholesterolemia (Garuti et al., 2005) and autosomal dominant polycystic kidney disease (Obermüller et al., 2001). These disparate clinical outcomes resulting from endocytic protein deficiency underscore the importance of investigations in the organismal context. Currently, many components of the endocytic machinery have been examined in only cell lines. In this study, our goal was to use an *in vivo* experimental system to investigate two important regulators of endocytosis, <u>PH</u> domain containing <u>e</u>ndocytic <u>t</u>rafficking <u>a</u>daptor 1 and 2 (PHETA1/2).

PHETA1 and PHETA2 (also known as Ses1/2 or IPIP27A/B or PHETA1/2) were identified *in vitro* as regulators of endosomal trafficking, specifically for receptor recycling to endosomes and for cargo sorting to lysosomes (Swan et al., 2010; Noakes et al., 2011). Both PHETA1 and PHETA2 have a C-terminal phenylalanine-histidine motif (F&H motif) that serves as a binding site for OCRL, encoded by a gene that is mutated in Lowe syndrome (MIM #309000)(Pirruccello et al., 2011). OCRL is an inositol 5-phosphatase, with phosphatidylinositol 4,5-bisphosphate (PI(4,5)*P_2_*) as the preferred substrate (Attree et al., 1992; Noakes et al., 2011). Binding to PI(4,5)*P_2_* occurs at the pleckstrin homology (PH) domain in OCRL, which also contains a loop outside the domain fold with a clathrin-binding motif. This motif directs OCRL specifically to clathrin-coated endocytic pits on the plasma membrane (Choudhury et al., 2009; Mao et al., 2009). PI(4,5)*P_2_* is abundant at the plasma membrane and is involved in a wide variety of processes, including actin dynamics and endocytosis (Sasaki et al., 2009). Disrupting OCRL’s phosphatase activity interferes with PI(4,5)*P_2_* homeostasis, which is thought to contribute to the disease manifestations of Lowe syndrome.

Several studies have shown that PHETA1 is critical in maintaining optimal OCRL function. Specifically, OCRL’s 5-phosphatase activity relies upon PHETA1-mediated interaction with PACSIN2 (<u>P</u>rotein kinase C <u>a</u>nd <u>c</u>asein kinase <u>s</u>ubstrate in <u>n</u>eurons 2), a protein that interacts with the actin cytoskeleton. A proline-rich PPPxPPRR motif in PHETA1 located upstream of the F&H motif serves as the necessary PACSIN2 binding site (Billcliff et al., 2016). PHETA2 lacks the PPPxPPRR motif). OCRL also promotes ciliogenesis by way of endosomal trafficking in a Rab8/PHETA1-dependent manner (Coon et al., 2012). These findings suggest that PHETA1 and OCRL functionally interact to mediate endocytosis and ciliogenesis.

Additionally, PHETA1 and PHETA2 are involved in the transport of newly synthesized lysosomal hydrolases from the trans-Golgi network (TGN) to the endosomes (Noakes et al., 2011). Thus, a loss of PHETA1/2 could result in improper sorting of lysosomal hydrolases. Consistent with this idea, loss of PHETA2 resulted in hypersecretion of pro-cathepsin D(Noakes et al., 2011). Similar disruptions in lysosomal proteins have also been found in mucolipidosis type II (MLII), where the loss of mannose 6-phosphate-dependent targeting resulted in hypersecretion of multiple lysosomal enzymes (Kudo et al., 2006; Koehne et al., 2016). Dysregulation of cathepsins in MLII zebrafish models resulted in craniofacial and skeletal deformations, recapitulating the clinical features of MLII patients (Spranger and Wiedemann, 1970; Cathey et al., 2010; Koehne et al., 2016). Thus, PHETA1/B dependent regulation of protease transport may be important for craniofacial development.

This hypothesis is supported by recent findings in a human patient with a *de novo* arginine to cysteine (R6C) variant in PHETA1, identified through the National Institute of Health’s Undiagnosed Diseases Program (UDP) (Gahl et al., 2012; Gahl et al., 2015; Gahl et al., 2016). This UDP patient presented with some clinical features of Lowe syndrome, but not the predominant manifestations of the disease, i.e., congenital cataracts, central nervous system abnormalities (congnitive impairment), and renal tubular and glomerular dysfunction (Mehta et al., 2014). These findings suggest that PHETA1 and OCRL may have both shared and independent functions *in vivo*.

To investigate the *in vivo* functions of PHETA1 and its close homolog PHETA2, we chose to utilize zebrafish, an informative small vertebrate model organism for validating the pathogenicity of genes or alleles in human patients. This approach has offered valuable insight for clinicians into a broad range of genetic disorders, including neurodevelopmental disorders, ciliopathies, and Lowe syndrome (Coon et al., 2012; Ramirez et al., 2012; Phillips and Westerfield, 2014; Oltrabella et al., 2015; Song et al., 2016; Sakai et al., 2018). Using histological, physiological, and behavioral analyses, we found that zebrafish *pheta1* and *pheta2* are required for endocytosis, ciliogenesis, and craniofacial development. The disruption in craniofacial development in the *pheta1/2* mutants was associated with a dysregulation in cathepsin K activity, likely due to its mislocalization. The abnormal craniofacial development is exacerbated further in the presence of the R6C variant, suggesting a dominant-negative mode of action in human disease.

## Results

### Identification of a *de novo* PHETA1 variant in undiagnosed human disease

The Undiagnosed Diseases Program (UDP) enrolled a 6-year old female patient with craniofacial dysmorphic features, scoliosis, clinodactyly, global developmental delay, vision and auditory impairments, and renal tubular or glomerular dysfunction (Fig. 1A-B and Table 1). Whole-exome sequencing and Sanger sequencing of the patient, unaffected fraternal twin, and unaffected parents identified a heterozygous *de novo* arginine (R) to cysteine (C) mutation in *PHETA1* (NM_001177997.2:c.55C>T; p.R6C in the short isoform, p.R19C in the long isoform, NM_001177996.1) only in the patient (Fig. 1C). The R6 residue in PHETA1 is highly conserved across species (Fig. 1D) (Papadopoulos and Agarwala, 2007), and the R6C mutation was predicted to be damaging with the use of Polyphen (Probably damaging, HumDiv: 1; HumVar: 0.995), SIFT (Deleterious, Score 0.01), and MutationTaster (Disease causing, Prob:1) (Adzhubei et al., 2010; Sim et al., 2012; Schwarz et al., 2014). This variant has been reported in ExAC browser with minor allele frequency of 0.000009398 (1/106410). Using patient-derived fibroblasts, we find that the R6C mutation does not affect the mRNA expression of *PHETA1* (Fig.1E)

**Figure 1.**
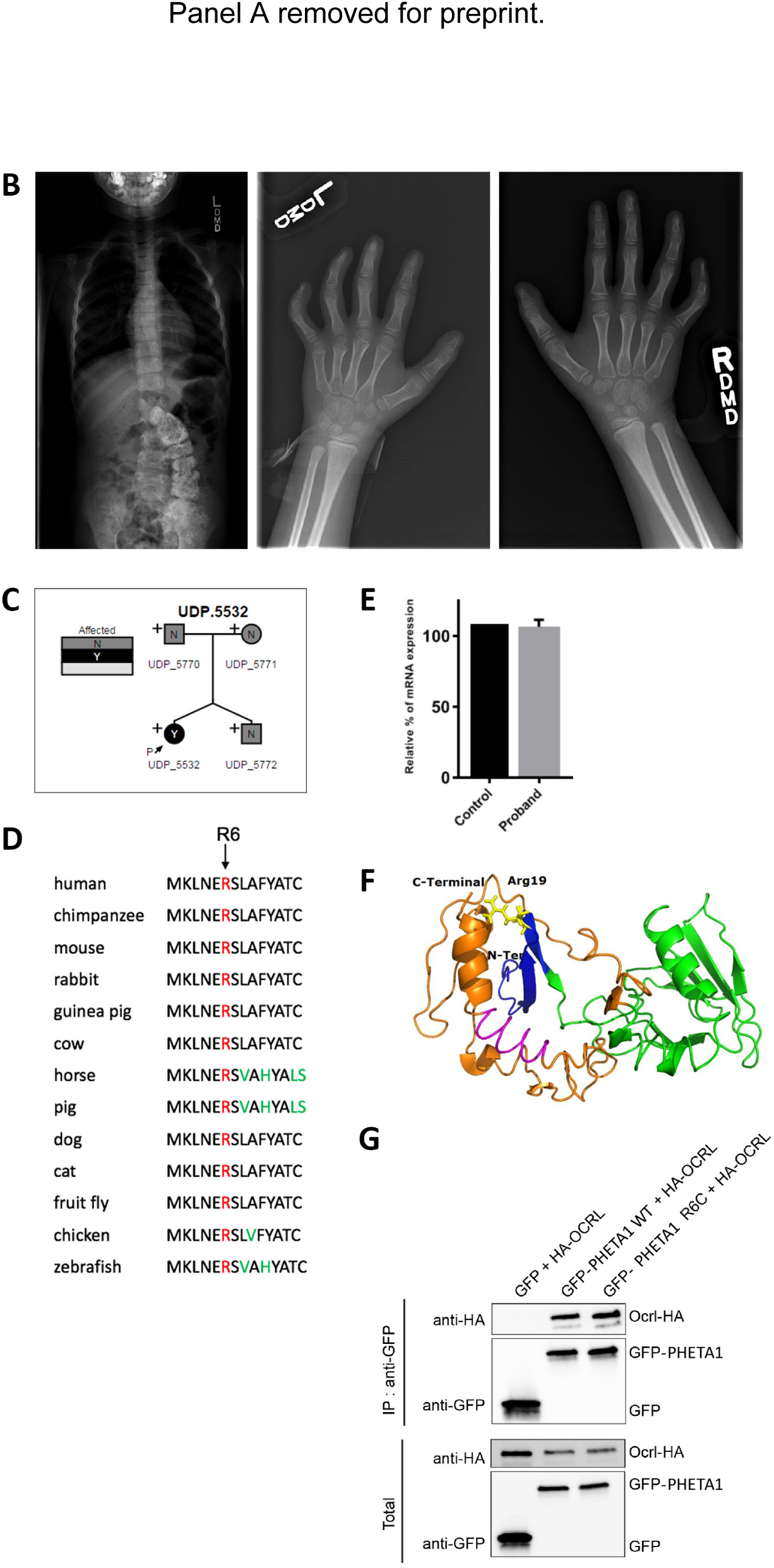
Identification of a *de novo* mutation in human PHETA1. (A) Images of UDP patient who presents with facial asymmetry, concave nasal ridge, and malar flattening. Radiograph reveals mild asymmetry of the skull. (B) Radiographs depict scoliosis and clinodactyly of fourth and fifth digits on both hands. (C) Whole exome sequencing was performed on both parents and fraternal twin of the UDP patient. N denotes “Not affected” and Y denotes “Affected.” P with the arrow identifies the UDP patient (UDP_5532). + indicates the presence of a normal allele, thus marking p.R6C as a heterozygous mutation. (D) COBALT multiple alignment of partial protein sequences of PHETA1 orthologs. The conserved arginine residue is highlighted in red, and amino acid residues that differ from the sequence of the human PHETA1 protein are highlighted in green. The arginine residue is highly conserved across multiple species. (E) Relative quantification of mRNA expression in the patient cells showing the expression of *PHETA1* is not significantly different compared to controls. Error bar represents standard deviation from six replicates. (F) 3-D structure of the human PHETA1 protein showing the PH domain (green) with a four-stranded N-terminal and three-stranded C- terminal β-sheet with a helix (orange). The conserved arginine amino acid (Arg19 in the PHETA1 long isoform, yellow) is far from the F&H motif (magenta); however, it stabilizes the folded domain around the C-terminal helix. (G) GFP-tagged full length WT PHETA1 or PHETA1^R6C^ were expressed in HeLa cells and tested for interaction with full length HA-tagged OCRL1. Bound proteins detected by Western blotting with indicated antibodies.

**Table 1.** Clinical features of UDP patient affected with a de novo arginine (R) to cysteine (C) mutation in PHETA1.

| Table 1. UDP_5532 Patient Clinical Presentations |  |
| --- | --- |
| Presentation at Birth | Abnormality of the Placenta, Breech presentation (C-section performed), Gestational diabetes, Congenital bilateral hip dislocation, Congenital muscular torticollis, Decreased body weight, Birth length less than 3<sup>rd</sup> percentile, Episodic vomiting, Feeding difficulties in infancy, Self-injurious behavior, Small for gestational age |
| Facial Features | Facial asymmetry, coarse facial features, concave nasal ridge, flat occiput, malar flattening, narrow mouth, sparse scalp hair, relative macrocephaly |
| Brain | Subdural hemorrhage, Absence of acoustic reflex, Ventriculomegaly, Elevated brain lactate level by MRS, Reduced brain N-acetyl aspartate level by MRS, Cavum septum pellucidum, Widened subarachnoid space, Decreased sensation to painful stimuli |
| Eyes | Congenital exotropia, amblyopia, bilateral ptosis, hypertelorism, nystagmus, optic nerve dysplasia/hypoplasia, short palpebral fissure, telecanthus. Patient had corrective surgery and vision is now normal. |
| Kidney | Horseshoe kidney, oligosacchariduria |
| Motor | Broad-based gait, delayed fine and gross motor development, generalized hypotonia, oral motor hypotonia |
| Hands/Feet | Clinodactyly of 4<sup>th</sup> and 5<sup>th</sup> finger, multiple palmar and plantar creases, pes planus, short foot and palm, tapered fingers, slow-growing nails, metatarsus adductus |
| Dental | Abnormality of dental morphology, dental malocclusion, difficulty in tongue movements, widely spaced teeth |
| Other | Moderate receptive language delay, severe global developmental delay, hearing impairment, thoracolumbar kyphoscoliosis, chronic constipation, mitral valve prolapse, freckled genitalia, hyperpigmented streaks, prominent crus of helix, upper airway obstruction, hip dysplasia, seasonal allergy |

Protein modeling was performed using the I-TASSER, MUSTER, and PHYRE2 servers (Fig. 1F) (Wu and Zhang, 2008; Adzhubei et al., 2010; Roy et al., 2010; Kelley et al., 2015). Based on the homology model, the arginine residue (highlighted in yellow) is far from the OCRL binding site (highlighted in magenta). However, it stabilizes the folded domain around the C-terminal helix, close to the F&H motif, such that the mutation is predicted to disrupt the folded domain and thus may interfere with OCRL binding to PHETA1. To test this, we expressed wild-type (GFP-PHETA1^WT^) and mutant (GFP-PHETA1^R6C^) GFP-tagged PHETA1 in HeLa cells, along with hemagglutinin (HA)-tagged OCRL. Surprisingly, we found that HA-OCRL was co-immunoprecipitated by both wild-type and mutant PHETA1 (Fig. 1G). This suggests that the R6C mutation might disrupt PHETA1 protein function in a manner that does not affect OCRL binding. Since the *in vivo* functions of PHETA1 and its close homolog, PHETA2, were still unknown, we used zebrafish as the experimental system to determine PHETA1/2’s roles in the context of an vertebrate organism.

### Expression and gene targeting of *pheta1* and *pheta2*

Like human, zebrafish has two PHETA proteins, which we will refer to as Pheta1 (encoded by *si:ch211-193c2.2*) and Pheta2 (encoded by *zgc:153733*). All known protein domain structures are conserved between human and zebrafish proteins, including the F&H motif (site of OCRL binding) (Fig. 2A). Pheta1 (but not Pheta2) contains the PPPxPPRR motif for PACSIN2 binding, like that of the human PHETA1. The neighboring genes of the human *PHETA1* and zebrafish *pheta1* loci are also conserved, suggesting that zebrafish *pheta1* is the most likely ortholog of *PHETA1* (Fig. S1A). The *pheta2* locus lacked obvious synteny with either the *PHETA1* or *PHETA2* loci. Based on overall amino acid sequence similarity, zebrafish Pheta1 is more similar to mammalian PHETA1 and PHETA2 (57.3% and 48.9% similarity, respectively), while Pheta2 is more divergent (44.2% and 39.2% similarity, respectively) (Fig. S1B).

**Figure 2.**
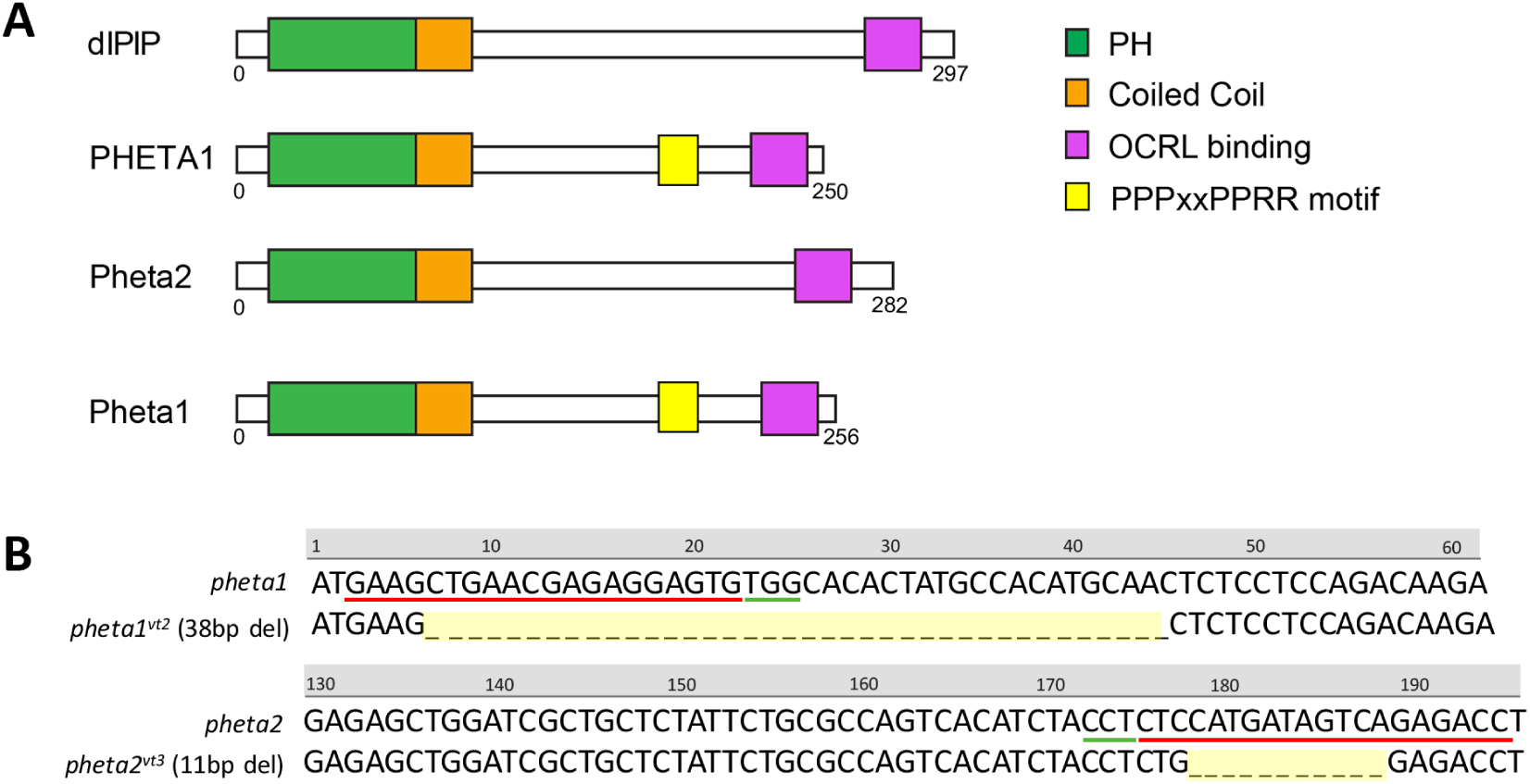
Pheta1 domain structure and CRISPR gene targeting. (A) The domain structures of *Drosophila melanogaster* homolog of PHETA (dIPIP), human PHETA1, and the zebrafish orthologs, Pheta1 and Pheta2. The PH domain, coiled-coil domain, OCRL binding site, and PPPxPPRR motif are highlighted. Like human PHETA2, zebrafish Pheta2 lacks the PPPxPPRR motif. (B) CRISPR mutagenesis. Top panel shows *pheta1* WT sequence and *pheta1^vt2^* sequence with 38bp deletion. Bottom panel shows *pheta2* WT sequence and *pheta2^vt3^* sequence with 11 bp deletion. CRISPR target site is underlined in red, and PAM site in green. Note that *pheta2* sequence shown is reverse complement.

To determine the functions of *pheta1 in vivo*, we generated a mutant allele of *pheta1* utilizing CRISPR genome engineering (Jao et al., 2013; Auer et al., 2014; Gagnon et al., 2014; Irion et al., 2014; Hisano et al., 2015). The resulting mutant allele, *pheta1^vt2^*, contains a thirty-eight base pair (38bp) deletion after the start codon, resulting in frame shift and predicted premature translational termination, suggesting that this is likely a null allele (Fig. 2B). In addition, in order to test for potentially redundant functions of *pheta1* and *pheta2*, we also generated a *pheta2* mutant allele. This mutant allele, *pheta2^vt3^*, contains an eleven base pair (11bp) deletion in exon 2, also resulting in a frame-shift and predicted premature translational termination, suggesting that this is also a null allele (Fig. 2B).

Zygotic *pheta1^vt2^* homozygous mutants (*pheta1-/-*) were viable and fertile, with no gross abnormalities during development. The same was true for zygotic *pheta2^vt3^* homozygous mutant animals (*pheta2-/-*). The presence of both the *pheta1* and *pheta2* transcripts in the 1-cell stage embryo indicated that this gene is maternally inherited (Fig. S2), and the maternal transcript might compensate for the loss of zygotic transcripts during the early stages of development. We therefore focused on the maternal-zygotic *pheta1* and *pheta2* mutants (progeny from homozygous mutant mothers), which lacked functional maternal and zygotic transcript during development. We also generated maternal-zygotic *pheta1-/-;pheta2-/-* double-knockout mutants (referred to as *dKO)* to test for redundant functions between the two PHETA proteins in the zebrafish. We found that maternal-zygotic *pheta1-/-*, *pheta2-/-*, and *dKO* animals were viable and fertile as well, with no gross abnormalities during development. Therefore, *pheta1* and *pheta2* were not required for viability and fertility in zebrafish.

### Loss of *pheta1* and *pheta2* impaired fluid-phase endocytosis

Previous findings in *ocrl* deficient zebrafish and the UDP patient phenotypes suggested that *pheta1/2* might regulate endocytosis in the renal system (Oltrabella et al., 2015). We examined renal endocytosis in the zebrafish, utilizing an established assay in which fluorescent tracers were injected into the common cardinal vein (CCV), followed by filtration and reabsorption into the renal tubular cells lining the pronephric kidney, commonly referred to as the pronephros (Drummond et al., 1998; Oltrabella et al., 2015). Endocytic uptake into the renal tubular cells can then be analyzed using fluorescent microscopy. We first tested fluid-phase endocytosis and micropinocytosis, using 10 kDa fluorescent dextran as the tracer (Li et al., 2015). Animals were injected at 3 days post fertilization (dpf) and then categorized into three groups: good endocytic uptake, low uptake, or no uptake (Fig. 3A). The *pheta1* heterozygous (*pheta1*^+/-^) and homozygous (*pheta1*^-/-^) animals showed a trend of reduced tracer uptake, compared to the wild-type (WT) control animals, but the difference did not reach statistical significance. The *dKO* animals, however, exhibited a significant reduction in tracer uptake compared to WT controls (Fig. 3B). This suggests that *pheta1* and *pheta2* acted redundantly for fluid-phase endocytosis, such that endocytic deficit was only observed when both proteins were depleted.

**Figure 3.**
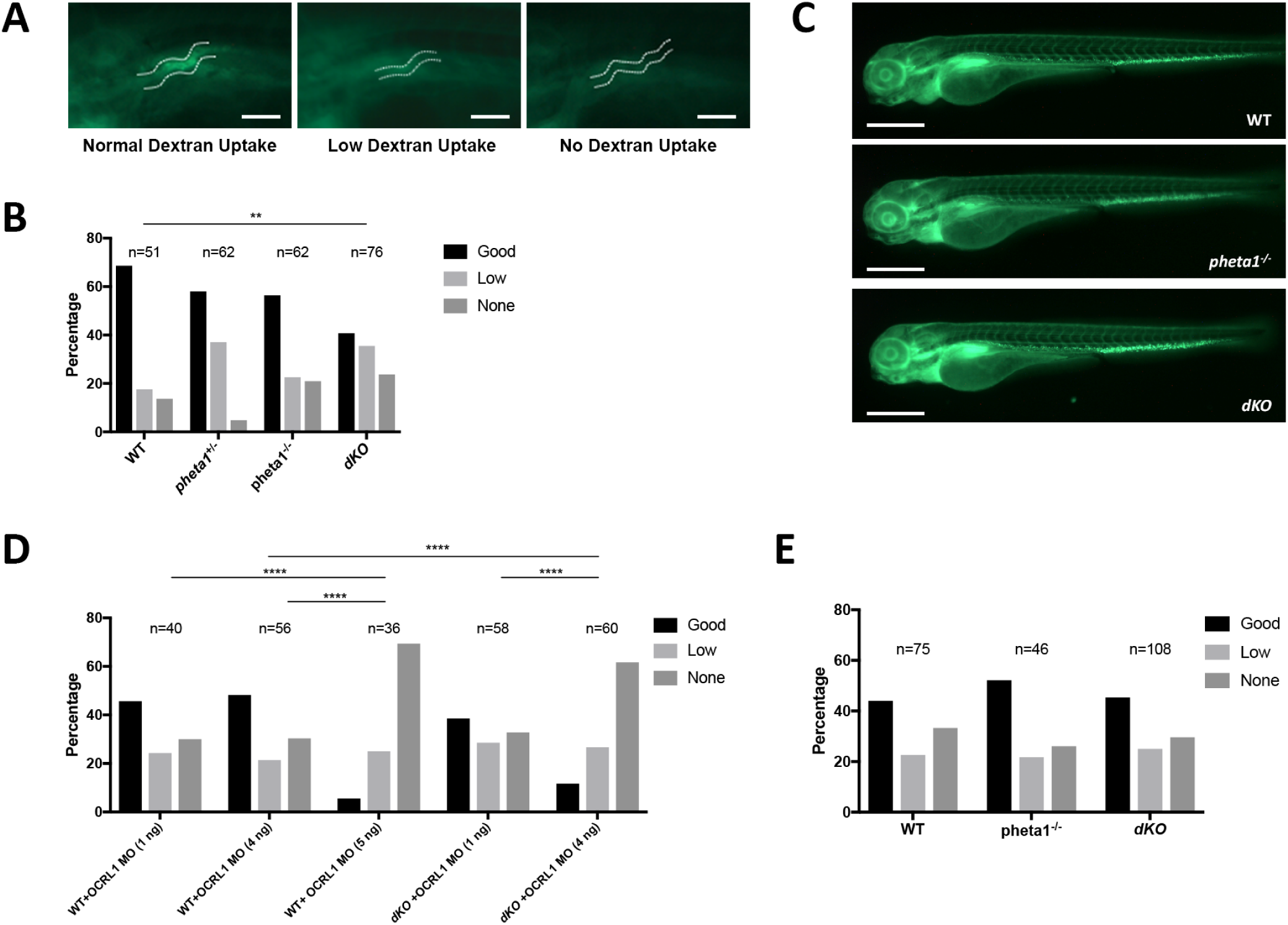
Loss of *pheta1/2* disrupts fluid-phase endocytosis. (A) Example fluorescent images of WT animals with good, low, or no dextran uptake. Scale bar= 100 µm. (B) Pronephros uptake of Alexa 488-10kDa Dextran. (C) Representative images of 3 dpf animals injected with 500 kDa Dextran, and imaged 24 hpi. Scale bar= 200 µm. (D) WT and *dKO* animals injected with *ocrl* MO at 1-cell stage, then 10 kDa dextran at 3 dpf. (E) Injection with RAP-Cy3. Statistical analyses performed using the Pearson’s chi-squared test. **p<0.01, ****p<0.0001. Abbreviations: WT (wild-type), MO (morpholino).

To verify that the reduction of 10 kDa dextran uptake in the *dKO* animals was a fluid-phase endocytosis-specific defect, and not due to the disruption of the glomerular filtration barrier, we tested glomerular filtration in the *pheta1-/-* and *dKO* animals by injecting 500 kDa dextran. 500 kDa dextran is too large to pass through a normally functioning glomerular filtration barrier, so it is expected to remain in the bloodstream. As shown in Fig. 3C, the 500 kDa dextran was retained in the bloodstream in both the *pheta1*-/- and *dKO* animals at 24 hours post-injection (hpi), like that of the WT controls. Together, these results show that *pheta1/2* are required specifically for fluid-phase endocytosis in the renal organ.

Consistent with the *in vitro* findings that OCRL and PHETA1/2 physically and functionally interact during endosomal trafficking, we found that reduction of zebrafish *ocrl* gene function significantly enhanced the fluid-phase endocytosis deficits in *dKO* animals. To inhibit *ocrl* function, we injected a previously validated *ocrl* anti-sense morpholino (MO) at the 1-cell embryonic stage and then later injected the 10kDa dextran (Coon et al., 2012). Injection of 5 ng/nl *ocrl* MO in the WT animals resulted in a severe reduction of dextran uptake, as previously described (Oltrabella et al., 2015). Injection of 4 ng/nl *ocrl* MO in WT animals results in a partial reduction of dextran uptake. The same *ocrl* MO concentration, however, resulted in a more severe reduction in dextran uptake in *dKO* animals (Fig. 3D). This suggests that *ocrl* and *pheta1/2* functionally interact in zebrafish to enable fluid-phase endocytosis. Interestingly, although *ocrl* is required for receptor mediated uptake of the receptor-associated protein (RAP) (Anzenberger et al., 2006; Oltrabella et al., 2015), we found no significant differences among WT, *pheta1-/-*, and *dKO* animals in RAP endocytic uptake (Fig. 3E). Together, these results indicate that *pheta1/2* are only involved in a subset of *ocrl*’s edocytic functions, specifically fluid-phase endocytosis.

### Loss of *pheta1/2* disrupted ciliogenesis in the pronephros

Several *in vitro* studies have described defects in ciliogenesis after OCRL depletion, and it was suggested that OCRL regulates protein trafficking to the cilia in a Rab8/PHETA1-dependent manner (Coon et al., 2012; Luo et al., 2012; Rbaibi et al., 2012). To determine if depletion of *pheta1* and/or *pheta2* affect ciliogenesis or cilia maintenance *in vivo*, we analyzed the cilia in the pronephros of 3 dpf larvae. We found that *dKO* animals had shorter and fewer cilia, similar to the phenotype seen in *ocrl*-deficient fish (Fig. 4) (Oltrabella et al., 2015). Since *dKO* animals exhibited a similar cilia phenotype like that of the *ocrl* mutants, this further supports the hypothesis that OCRL and PHETA proteins are involved in the same pathway.

**Figure 4.**
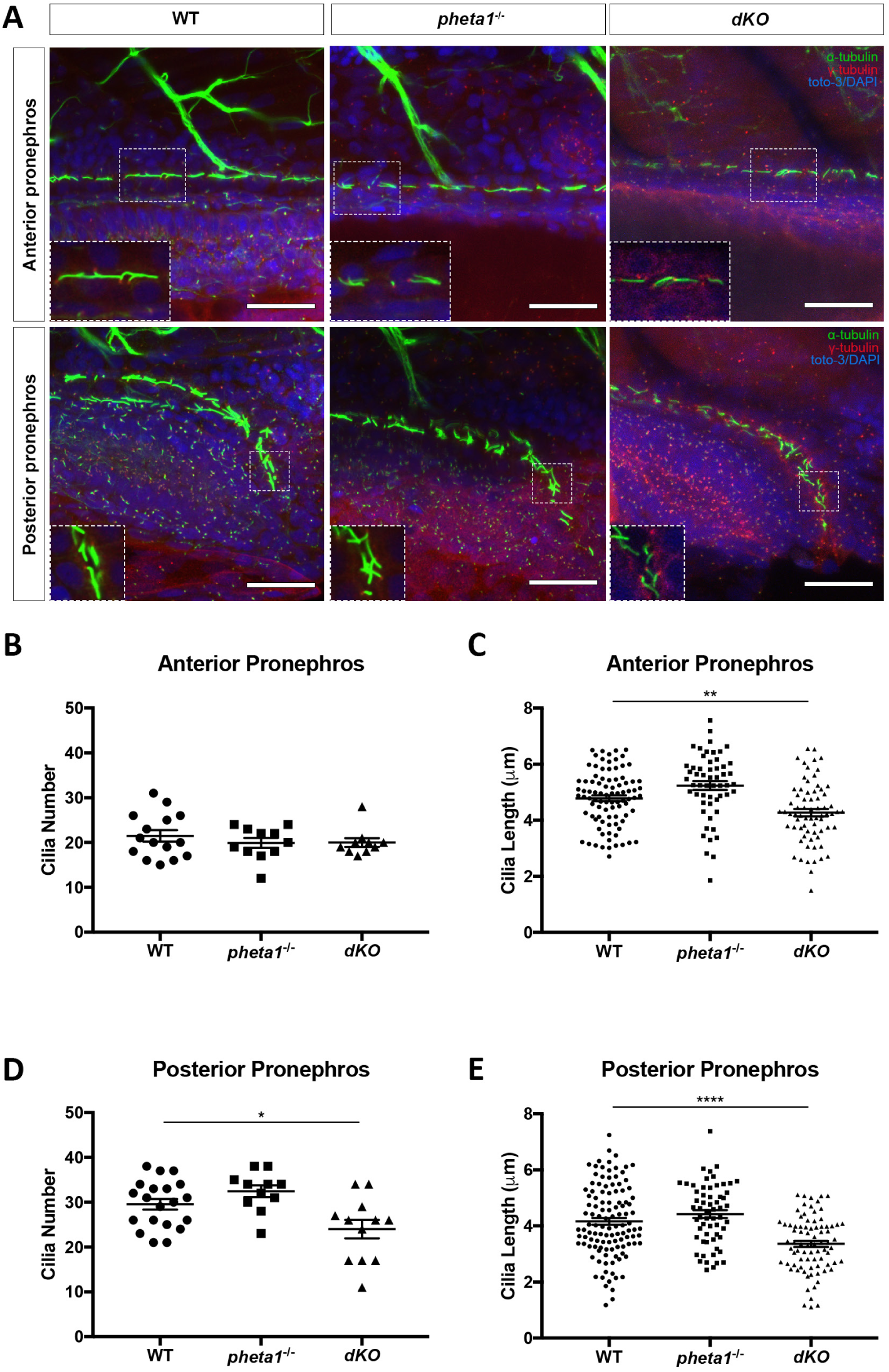
Loss of *pheta1/2* disrupts ciliogenesis in the pronephros. (A) Representative confocal images of cilia in the pronephros of WT, *pheta1*^-/-^, and *dKO*. Cilia were labeled with anti-acetylated α-tubulin (green), basal bodies labeled with anti-γ tubulin (red), and nuclei labeled with toto-3 or DAPI (blue). Scale bar= 25 µm. Insert depicts magnified image of individual cilia. (B) Cilia number in anterior pronephros. WT: n=15, *pheta1*^-/-^: n=11, *dKO*: n=10. (C) Cilia length in anterior pronephros. WT: n=15, *pheta1*^-/-^: n=11, *dKO*: n=10. Five cilia selected from each animal for cilia length measurements. (D) Cilia number in posterior pronephros. WT: n=20, *pheta1*^-/-^: n=11, *dKO*: n=12. (E) Cilia length in posterior pronephros. WT: n=20, *pheta1*^-/-^: n=11, *dKO*: n=12. Five cilia selected from each animal for cilia length measurements. Error=SEM. Statistical analyses performed using one-way ANOVA with Holm-Sidak’s multiple comparisons test. *p<0.05, **p<0.01, ****p<0.0001.

We next examined if there was a potential ciliary deficit in other ciliated organs, including the inner ear (macula and crista), the olfactory placode, and the lateral line. We found no disruption of ciliogenesis or cilia maintenance in these tissues analyzed in *pheta1-/-* and *dKO* animals (Fig. S3). We also examined the outer segments of photoreceptors, which are specialized cilia that originate from the apical-most region of the inner segment. Staining with rod and cone photoreceptor markers did not reveal any differences among WT, *pheta1-/-*, *pheta2-/-*, and *dKO* animals (Fig. S3). This suggests that *pheta1/2*’s role in ciliogenesis was restricted to the pronephros.

### *pheta1/2* is not required for oculomotor function

The UDP patient was born with multiple visual complications, including deficits with oculomotor function (e.g., congenital exotropia, amblyopia, nystagmus). To test whether a deficiency in PHETA1/2 contributes to oculomotor deficits, we analyzed the optokinetic response (OKR) in the *pheta1/2* mutants. OKR is a gaze stabilization response that utilizes the extraocular muscles to stabilize a image on the retina in response to visual motion. It is necessary for maintaining optimal visual acuity and is conserved in all vertebrates (Huang and Neuhauss, 2008). In zebrafish, OKR develops early and is robust by 3-4 dpf (Easter and Nicola, 1997; Huang and Neuhauss, 2008). To ensure complete development of OKR, we tested 5-6 dpf larvae. Animals were placed inside a circular arena with projections of moving black and white gratings, and the eye positions were video-recorded (Fig. S4A). The grating directions alternated clockwise and counter-clockwise at different contrasts at 3-second periodicity. We tested siblings in the progeny of *pheta1/2* double heterozygous animals (*pheta1+/-;pheta2+/-*) and the eye velocity in response to the moving gradients was analyzed. No significant differences in eye velocity were observed among WT, *pheta1-/-*, *pheta2-/-*, or *dKO* siblings. We then investigated the correlation of velocity and angle between the left and right eyes (Fig. S4B-C). A reduced correlation would suggest strabismus, in which the eyes do not properly align with each other. No significant differences in angle or velocity correlation were observed among WT, *pheta1-/-*, *pheta2-/-*, or *dKO* siblings. Thus, the deficiency in *pheta1/2* did not affect oculomotor function.

### Loss of *pheta1/2* disrupts craniofacial morphogenesis

As mentioned previously, PHETA1 and PHETA2 were previously found to be involved in the sorting of lysosomal hydrolases *in vitro*. Disruption of this pathway *in vivo* could give rise to craniofacial abnormalities as seen in lysosomal storage disorders such as MLII (Kudo et al., 2006; Koehne et al., 2016). We investigated craniofacial development in our *pheta1/2* mutants by performing morphometric measurements of cartilage structures in 6 dpf larvae stained with Alcian Blue (cartilage) and Alizarin Red (bone). WT, *pheta1-/-*, *pheta2-/-*, and *dKO* animals were analyzed. Representative lower jaw (Fig. 5A, 6A) and upper jaw (neurocranium, Fig. 5B) images of the WT and mutants are shown. We did not observe any loss of craniofacial structures, but the mutants tended to have shorter and narrower jaws. To quantitate this potential difference, we made three primary measurements: a) cranial distance; b) jaw width; c) ceratohyal length (Fig. 5C-F). Together, these measurements informed us about the overall length and proportions of the head, as well as the growth of an individual cartilage structure (ceratohyal). Indeed, we found a significant reduction in these three metrics in the *pheta2-/-* and *dKO* animals, while the loss of *pheta1-/-* alone only mildly (though significantly) affected jaw width. A similar trend was seen in other craniofacial measurements (Fig. S5). Depletion of both *pheta1* and *pheta2* had an additive effect, indicating functional redundancy during craniofacial development.

**Figure 5.**
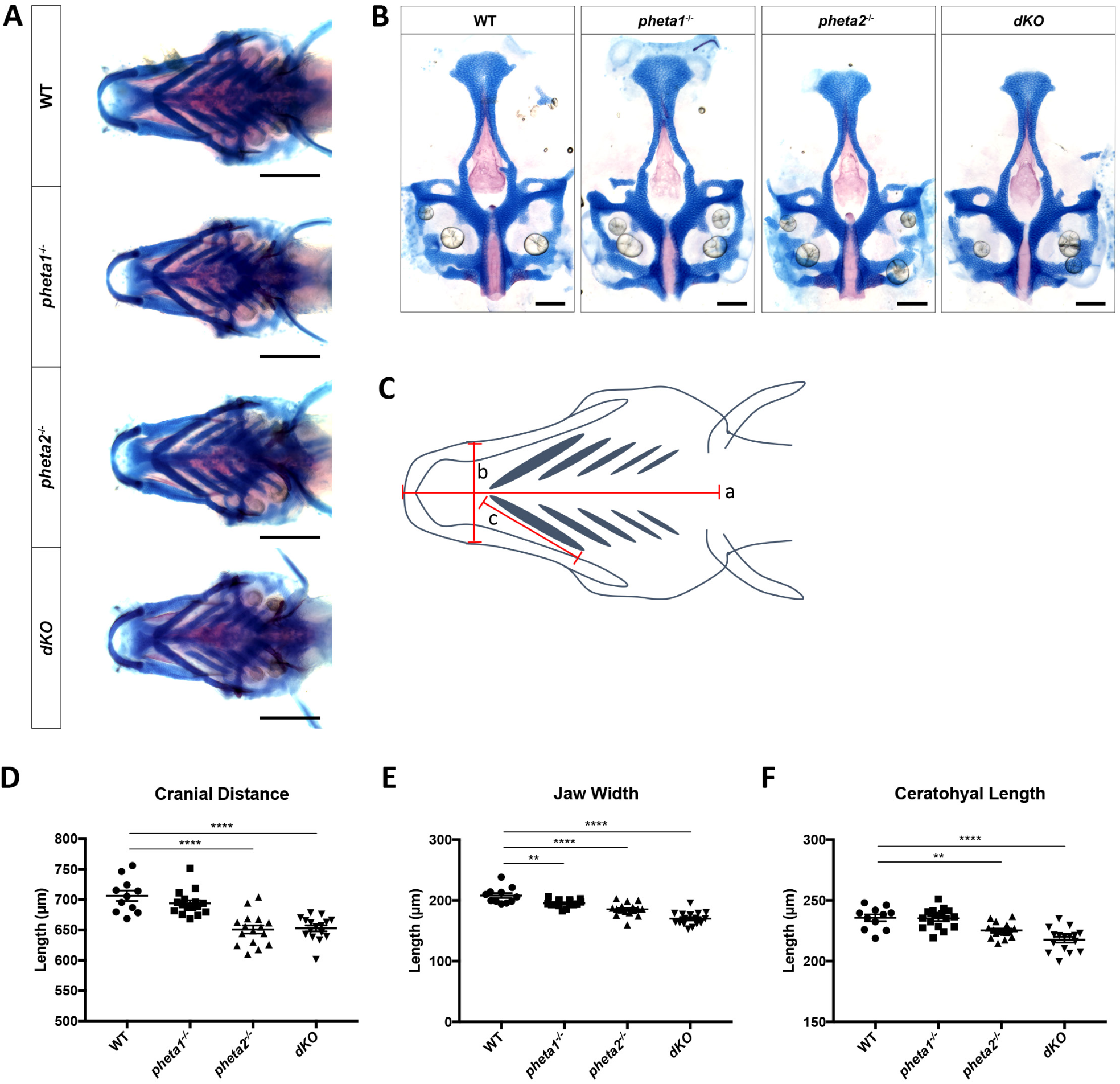
Loss of *pheta1/2* disrupts craniofacial development. (A) Representative images from each genotype analyzed. Scale bars= 200 µm. (B) Flat-mount preparations of neurocranium. Scale bars=100 µm. (C) Schematic with measurements analyzed: (a) cranial distance, (b) jaw width, and (c) ceratohyal length. (D-F) Analyses of craniofacial measurements, including cranial distance, jaw width, and ceratohyal length. Statistical analyses performed using one-way ANOVA with Holm Sidak post-test. WT: n=11, *pheta1^-/-^*: n=16, *pheta2^-/-^*: n=16, *dKO*: n=16. **p<0.01, ****p<0.0001.

**Figure 6.**
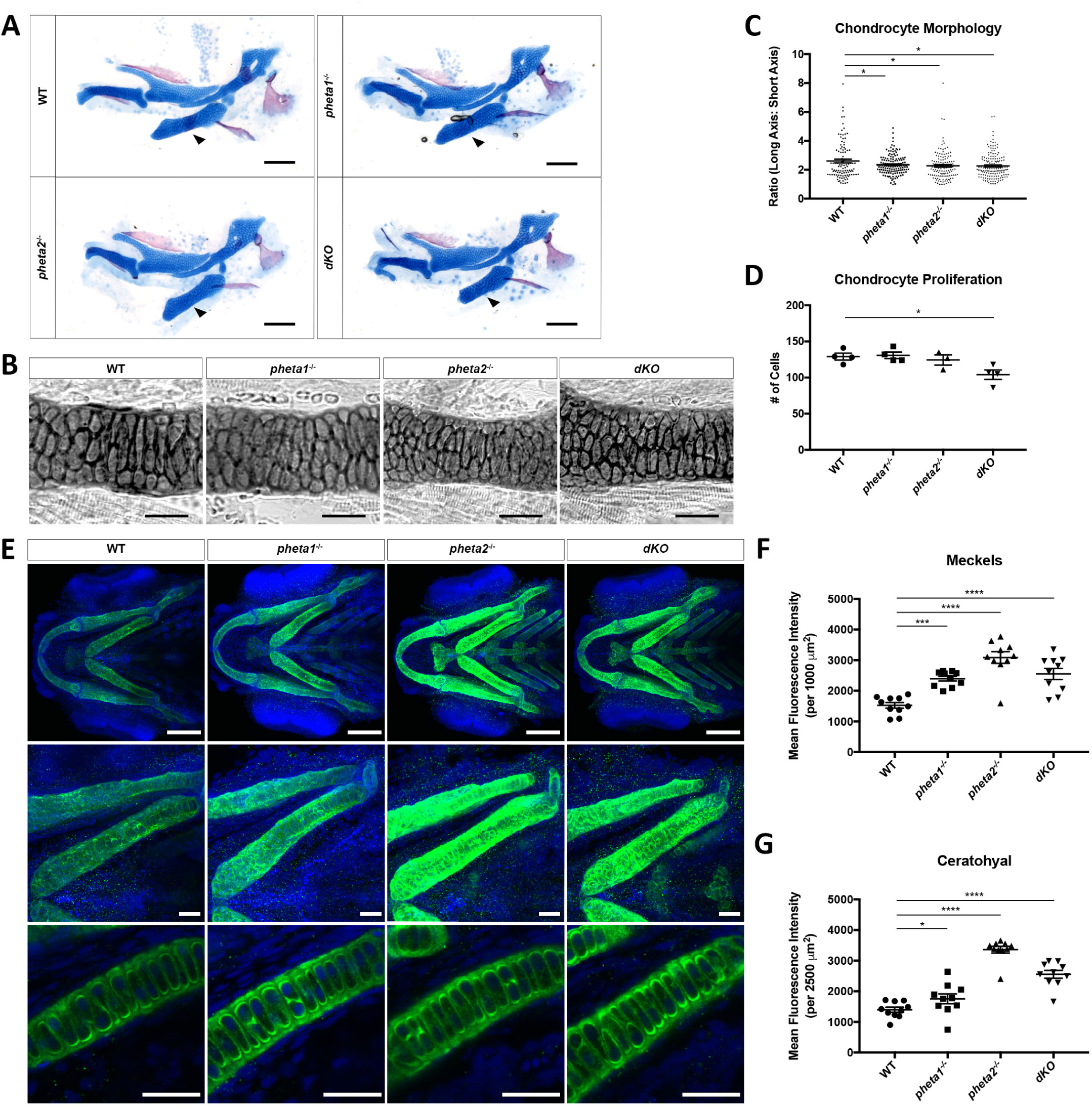
Depletion of *pheta1/2* disrupts chondrocyte proliferation and maturation. (A) Flat-mount preparations of the jaw. Arrowheads point to ceratohyal cartilage. Scale bar= 100 µm. (B) Ceratohyal cartilage at higher magnification. Scale bar= 50 µm. (C) Chondrocyte morphology analysis in ceratohyal cartilage (within 200 µm^2^ area). (D) Chondrocyte number in ceratohyal cartilage. Statistical analyses performed using one-way ANOVA with Holm Sidak post-test. n=3-4 animals per genotype. *p<0.05. (E) Top panel: Ventral view of 5 dpf larvae immunostained for type II collagen (green). Nuclei are labeled with DAPI (blue). Scale bar= 100 µm. Middle panel: Higher magnification of corresponding ceratohyal cartilage. Scale bar= 25 µm. Bottom panel: Higher magnification of corresponding ceratohyal cartilage; one optical section depicting extracellular secretion of type II collagen. Scale bar= 25 µm. (F) Quantification of mean fluorescence intensity in the Meckel’s cartilage. (G) Quantification of mean fluorescence intensity in the ceratohyal cartilage. Statistical analyses performed using one-way ANOVA with Holm Sidak post-test. n=10 animals per genotype. *p<0.05, ***p<0.001, ****p<0.0001.

### Loss of *pheta1/2* disrupts chondrocyte maturation

The craniofacial morphogenesis defects observed in *pheta1/2* mutants suggest an underlying developmental abnormality. To test this, we first examined the morphology of chondrocytes, which assume a more elongated shape when mature (Goldring et al., 2006). If there is a delay in early chondrocyte maturation, the chondrocytes can persist with more rounded morphology, with a long axis:short axis ratio closer to one. This can be measured in flat-mount preparations of the lower jaw (Fig. 6A-B). We examined the chondrocytes in the ceratohyal cartilage and analyzed the ratio of the long axis to the short axis of individual cells. We found a slight but significant reduction in the elongation of chondrocytes in the *pheta1-/-*, *pheta2-/-*, and *dKO* animals, suggesting a delay in chondrocyte maturation (Fig. 6C). We then asked if differences in chondrocyte proliferation could account for the reduced length of cartilage structures. Indeed, there was a 20% reduction in the number of cells in the ceratohyal cartilage of *dKO* animals compared to WT controls (Fig. 6D). No significant differences in cell number were observed in the Meckel’s cartilage (data not shown). Taken together, the altered chondrocyte morphology and reduced chondrocyte proliferation seen in the ceratohyal cartilage suggest that there was a disruption in proper chondrocyte maturation and differentiation in the *pheta1/2* mutants.

Next, we examined the expression of various markers that characterize the sequential stages of chondrocyte maturation. During chondrocyte maturation, there is a decline in TGF-β-signaling, and thus a decrease in Smad2/3 and Sox9 transcriptional regulators. At the same time, there is a coordinated change in the extracellular matrix protein composition as the chondrocytes mature (Goldring et al., 2006; Flanagan-Steet et al., 2016). Type II collagen (Col2, encoded by *col2a1a*) is one of the earliest markers of chondrocyte differentiation whose expression is decreased at later stages of development (Goldring et al., 2006; Flanagan-Steet et al., 2016). Immunostaining for Col2 at 4 dpf showed that *pheta2-/-* and *dKO* animals exhibited a striking increase compared to WT controls, while the *pheta1-/-* animals had a modest increase (Fig. 6E-G). This is consistent with our morphometric analyses, in which *pheta2-/-* and *dKO* animals exhibited a more severe deficit in overall craniofacial development compared to *pheta1*-/- animals. Due to the apparent increase in protein expression of Col2, we performed quantitative RT-PCR to investigate RNA expression levels of *col2a1a* (Fig. S6A). We found no significant increase in *col2a1a* expression at 4dpf, suggesting that the increased Col2 protein levels may be a result of altered translation or matrix turn over. We also analyzed *acana* (aggrecan), a later-stage extracellular matrix marker, and found a trend for decreased *acana* expression in the *pheta2-/-* and *dKO* animals at 4 dpf, compared to WT (Fig. S6B). Lastly, we wanted to determine if there is a sustained upregulation in the *sox9a* transcription factor at later stages. The Sox9 transcription factor is required for early chondrocyte development, but sustained Sox9 expression can inhibit later stages of maturation. At 4 dpf, in situ hybridization revealed an increase in *sox9a* expression in portions of the lower jaw (indicated by black arrowheads) in the *pheta2-/-* and *dKO* animals (Fig. 7A). Together, these molecular profiling findings support the hypothesis that a delay in chondrocyte maturation results in a deficiency in craniofacial morphogenesis.

**Figure 7.**
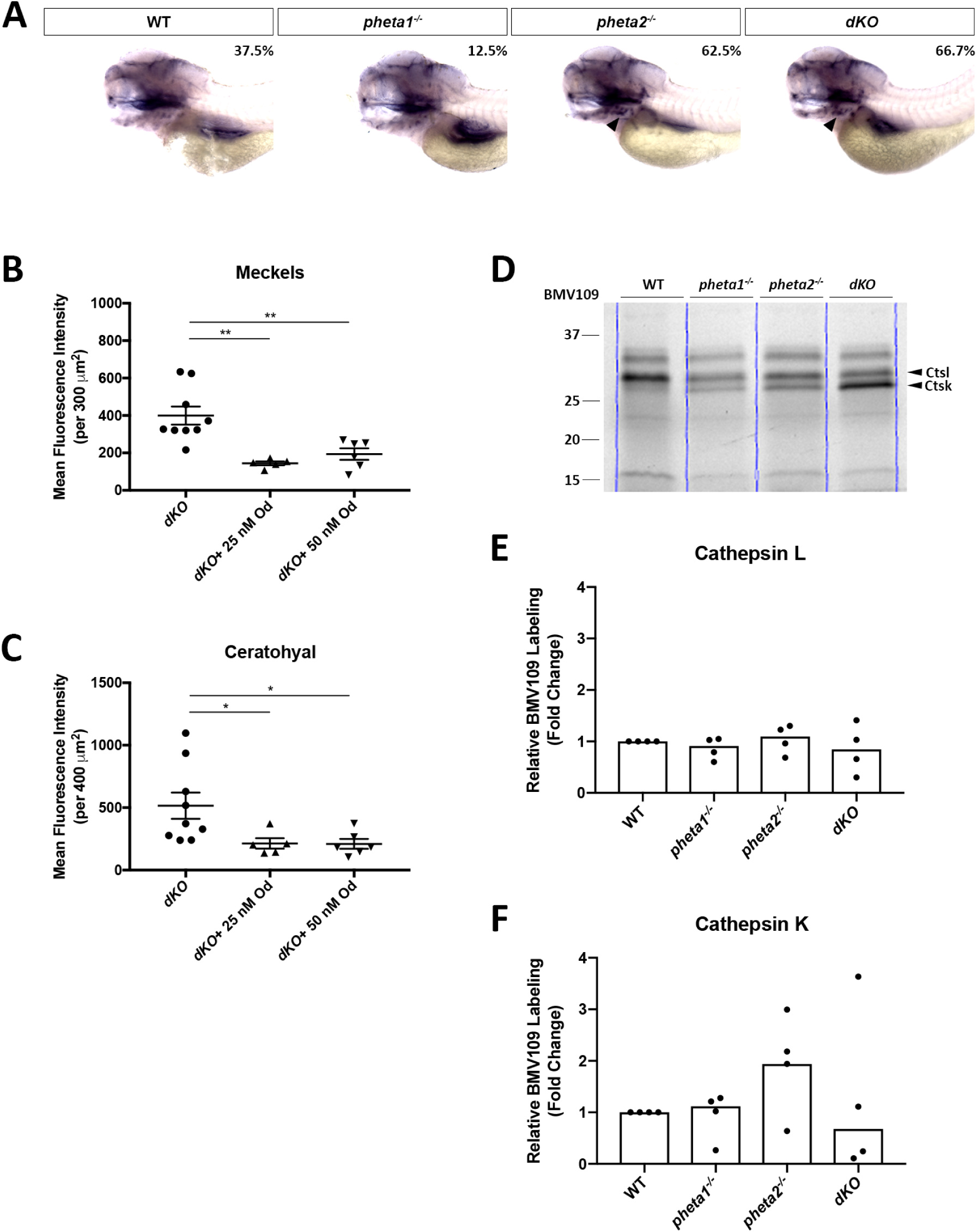
Loss of *pheta1/2* disrupts chondrocyte maturation and results in aberrant cathepsin K activity. (A) *In situ* hybridization of *sox9a* in WT and *pheta1/2* mutants. Arrowheads indicate regions of difference in lower jaw. Percentage with elevated lower jaw expression: WT=37.5%, n=8; *pheta1^-/-^*=12.5%, n=7; *pheta2^-/-^*=62.5%, n=8; *dKO* =66.7%, n=6. (B) Quantification of mean fluorescence intensity of type II collagen (*col2a1a*) in the Meckel’s cartilage after administration of odanacatib (Od). (C) Quantification of mean fluorescence intensity of type II collagen in the ceratohyal cartilage after administration of odanacatib (Od). (D) In-gel analyses of BMV109, showing cathepsin activities in WT and *pheta1/2* mutants at 4dpf. (E-F) Quantitation of the cathepsin K (Ctsk) and cathepsin L (Ctsl) bands from 4 experiments. Statistical analyses performed using one-way ANOVA with Holm Sidak post-test. Error=SEM. n=5-6 animals per genotype. *p<0.05, **p<0.01.

### *pheta1/2* regulates Col2 expression through cathepsin K

Extracellular matrix remodeling and homeostasis during chondrogenesis and osteogenesis are dependent on the function of proteolytic enzymes, including cysteine proteinases (known as cathepsins), and metalloproteinases (Yasuda et al., 2005; Holmbeck and Szabova, 2006). For example, dysregulation of cathepsin K causes craniofacial abnormalities in zebrafish models of the lysosomal storage disorder, MLII (Flanagan-Steet et al., 2018). Since PHETA1/2 was found to be involved in the transport of lysosomal hydrolases from the TGN to the endosomes, we hypothesized that cathepsin K dysregulation caused by a deficiency in *pheta1/2* may also cause deficits in craniofacial development. To test this, animals were treated with a cathepsin K-specific inhibitor, odanacatib, at 3 dpf and then collected and immunostained for Col2 at 4 dpf (Gauthier et al., 2008; Flanagan-Steet et al., 2018). Odanacatib significantly reduced Col2 in the *dKO* animals at both 25 nM and 50 nM concentrations (Fig. 7B-C). Interestingly, there was no reduction in *pheta1-/-* and *pheta2-/-*, indicating that *pheta1* and *pheta2* my be able to compensate for one another in regulating cathepsin K activity (Fig. S7).

Next, we asked if *pheta1/2* affects the level of cathepsin K activity. We utilized a cathepsin-specific activity-based probe (ABP), BMV109, to measure cathepsin activity in both WT and *pheta1/2* mutants (Verdoes et al., 2013; Flanagan-Steet et al., 2018). The animals were treated at 3 dpf, a period when several cathepsin activities typically begin to wane in WT animals (Flanagan-Steet et al., 2018). *pheta1/2* deficiency did not significantly impact cathepsin L activity. Cathepsin K activity, however, was more variable in the *pheta1/2* mutant animals, with a trend toward increased activity in *pheta2-/-* animals (Fig. 7D-F). Thus, the rescue effects of a cathepsin K inhibitor in *dKO* animals may reflect another mechanism of dysregulation or a broader imbalance of protease activity.

### *pheta1*^R6C^ exerts a dominant-negative effect on craniofacial development

One striking clinical feature identified in the UDP patient is her abnormal craniofacial development. The patient presented with coarse facial features and facial asymmetry. She also had shorter feet and palms, as well as dental abnormalities, including skeletal malocclusion. This could be due to *PHETA1* haploinsufficiency or dominant-negative effects of the R6C allele. If *pheta1*^R6C^ was non-functional or partly functional, then ectopic expression of Pheta1^R6C^ in *pheta1+/-* or *pheta1-/-* backgrounds should have no effect or partially improve craniofacial development. However, if *pheta1*^R6C^ was dominant-negative, then ectopic expression of *pheta1*^R6C^ should worsen craniofacial development in the same backgrounds. Thus, we generated two zebrafish transgenic lines, one that ubiquitously expressed an EGFP-Pheta1^R6C^ fusion protein [*Tg*(*R6C*)], and another that expressed EGFP fused with WT Pheta1 [*Tg(WT)*] (Fig. 8A). Confocal imaging confirmed the broad expression of *Tg*(*R6C*) and *Tg*(*WT*). (Fig. 8B).

**Figure 8.**
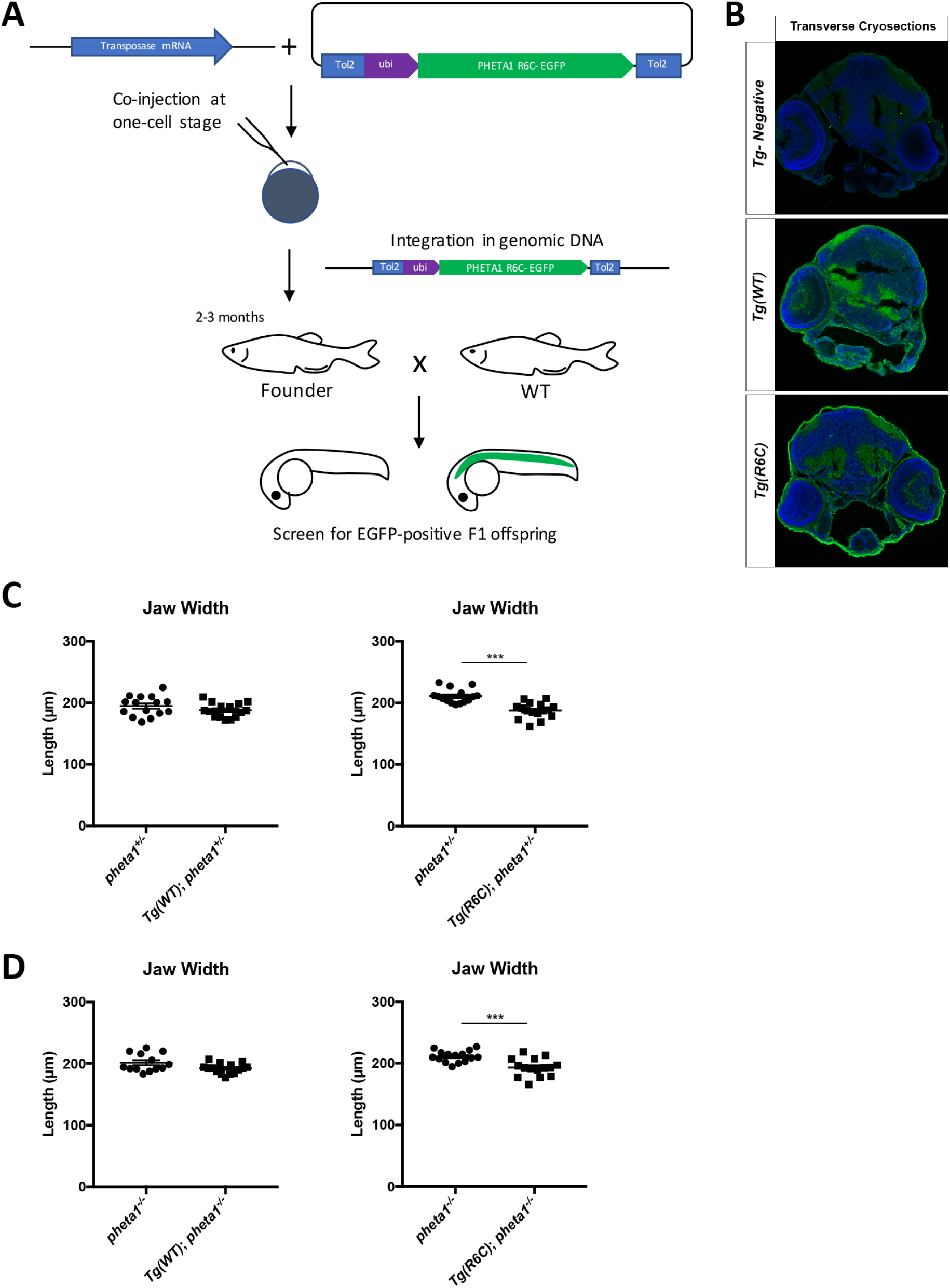
Pheta1^R6C^ exerts a dominant-negative effect on craniofacial development in the partial or complete absence of Pheta1. (A) Generation of *Tg(R6C)* and *Tg(WT)* transgenic lines. (B) Confocal images showing broad expression of Pheta1^WT^ and Pheta1^R6C^ in transverse cryosections of 3dpf larvae (indicated in green with anti-GFP). An EGFP-negative control is shown for comparison. DAPI was used as a nuclear stain (blue). (C) Jaw width in the *pheta1* heterozygous (*pheta1* ^+/-^) background. (D) Jaw width in the *pheta1* homozygous (*pheta1* ^-/-^) background. Statistical analyses performed using one-way ANOVA. n=15-16 animals per category. ***p<0.001, ****p<0.0001.

To mimic the genetic background of the UDP patient, which is heterozygous for the R6C allele, we analyzed the effects of *Tg(R6C)* and *Tg(WT)* in the *pheta1* heterozygous (*pheta1*^+/-^) background. We also analyzed the effects of *Tg(R6C)* and *Tg(WT)* in the *pheta1* homozygous (*pheta1*^-/-^) background to test whether there could be an effect in the absence of *pheta1*. We found that, in the partial and complete absence of *pheta1*, *pheta1*^R6C^ exacerbated the jaw width defect (Fig. 8C-D). This suggest that the R6C allele has dominant-negative effects. Transgenic expression of WT Pheta 1 had not significant effects in either *pheta1+/-* or *pheta1-/-* backgrounds, indicating that the effect on jaw width was specific to the expression of the R6C variant. There was no effect on cranial distance or ceratohyal length (Fig. S8). Interestingly, Pheta1^R6C^ impacted the craniofacial morphometrics that were affected by the loss of both *pheta1* and *pheta2* (jaw width), but not the morphometrics that were only affected by the loss of *pheta2* (cranial distance and ceratohyal length). This suggests that Pheta1^R6C^ may have a relatively limited capacity to interfere with Pheta2 function.

## Discussion

The regulation of endocytic trafficking is essential for the development and function of an organism. In this study, we present the first *in vivo* investigation of the functions of PHETA proteins, which are membrane adaptor proteins for the Lowe syndrome causative protein, OCRL. Using zebrafish as the experimental system, we found that *pheta1* and *pheta2* were necessary for renal fluid-phase endocytosis and ciliogenesis. Furthermore, we found that loss of *pheta1/2* impaired craniofacial development and altered the composition of the cartilage extracellular matrix. Evidence also indicates that cathepsin K is a contributing factor dysregulated by the deficiency of *pheta1/2* during craniofacial development.

These findings provide insight into the possible pathophysiology of an individual with a *de novo* R6C mutation in PHETA1. The patient presented with renal and craniofacial phenotypes that were similar to the observed phenotypes in *pheta1/2* mutant zebrafish, suggesting that deficiency in PHETA1 contributes to disease. Using transgenic expression in zebrafish, we found that the R6C allele acted in a dominant-negative manner. Together, we reveal the essential physiological and developmental roles of PHETA proteins and indicate cathepsin proteases as potential targets for PHETA-associated diseases.

### The roles of *pheta1* and *pheta2* in fluid-phase endocytosis and ciliogenesis

Loss of *pheta1/2* affected the renal fluid-phase endocytosis (of 10 kDa dextran substrate), but not receptor-mediated endocytosis (of RAP) (Anzenberger et al., 2006). In contrast, loss of *ocrl* in zebrafish resulted in a strong reduction in both types of endocytosis (Oltrabella et al., 2015). Partial knockdown of *ocrl* in the *dKO* animals exacerbated the fluid-phase endocytic deficit, indicating that *pheta1/2* and OCRL likely function in a common endocytic pathway. These results suggest that *pheta1*/*2* participates in only a subset of OCRL’s functions *in vivo*. Likely, other F&H motif-containing OCRL adaptor proteins such as APPL1 can partially compensate for the loss of PHETA1/2 (Swan et al., 2010; Noakes et al., 2011; Pirruccello et al., 2011).

The pronephros of *dKO* animals had fewer and shorter cilia, similar to what was found in *ocrl*-deficient zebrafish (Oltrabella et al., 2015). However, for several reasons, this mild ciliogenesis defect in both *pheta dKO* and *ocrl^-/-^* mutants cannot account for the observed impairment of endocytosis. First, unabsorbed fluorescent dextran was normally excreted from the cloacae in the *dKO* animals, indicating that there was no impairment of cilia-directed fluid flow within the pronephros. Second, we did not see the development of renal cysts in any of our *pheta1/2* mutants, which is consistent with normal fluid flow. Lastly, even mutants with severe ciliogenesis deficits (e.g., the *double bubble* mutant) could endocytose dextran normally (Drummond et al., 1998; Liu et al., 2007; Oltrabella et al., 2015). Thus, *pheta1/2* likely contributes to fluid-phase endocytosis independently of its role in ciliogenesis.

### A novel role for *pheta1* and *pheta2* in craniofacial development

We identified a novel role for *pheta1* and *pheta2* in craniofacial morphogenesis. Craniofacial development appeared to rely more on *pheta2*, but depletion of both *pheta1* and *pheta2* resulted in an additive effect, indicating that *pheta1* plays a role as well. In *pheta2-/-* and *dKO* animals, we observed features indicative of abnormal chondrocyte maturation, including abnormal chondrocyte morphology, changes in marker gene expression, and altered extracellular matrix composition (i.e., increased Col2). As a first foray into the underlying molecular mechanisms, we found that inhibition of cathepsin K using the specific inhibitor odanacatib significantly reduced Col2 protein levels in the *dKO* animals. This indicated that aberrant cathepsin K activity contributes to abnormal craniofacial development. Interestingly, we did not see a consistent systemic increase in cathepsin K activity using an in vivo activity probe (BMV109), indicating that the dysregulation of cathepsin K activity may be regional or stem from its mislocalization. Future studies might explore where active cathepsin K resides as development progresses and how Col2 levels are modulated by protease activity in *pheta1/2* mutant animals. Our previous findings have shown that TGF-β signaling is enhanced by cathepsin K activity, which may, in turn, mediate the abnormal craniofacial morphogenesis in the absence of *pheta1/2*.

### Investigating the pathogenesis of the UDP patient’s disease

A primary motivation for understanding the in vivo function of PHETA1/2 was the identification of a patient carrying a *de novo* PHETA1 mutation. While the R6C mutation did not affect interaction with OCRL, it did exert a dominant-negative effect on craniofacial development, even in the absence of endogenous *pheta1*. Since the R6C mutant can interact with OCRL, it may be able to compete with other OCRL interacting protein such as PHETA2 or APPL1. Alternatively, since PHETA1 and PHETA2 can form homodimers and heterodimers, the R6C mutant may bind to and interfere with the normal functions of PHETA1 and PHETA2. Our hypothesis that the R6C mutation resulted in a deficiency of PHETA1/2 function is supported by the overlapping phenotypes between the patient and our zebrafish mutants, specifically in craniofacial development and renal function (Table 1).

We note that the UDP patient has three other *de novo* mutations considered less likely to be contributing to disease. One variant in *DNAJB5* (DnaJ heat shock protein family (Hsp40) member B5; NM_001135004: p.R419H) has inconsistent predictions with SIFT and Polyphen, and occurs in a moderately conserved amino acid, so it is unlikely that this causes the UDP patient’s disease. A second variant, in *UPP1* (Uridine phosphorylase 1; NM_003364:p.I117V), is seen in 12 normal individuals and is predicted benign by SIFT and Polyphen, so it is unlikely to be pathogenic. The third variant, is in *PHF6* (Plant homeodomain (PHD)-like finger protein 6; NM_001015877.1: p.Leu244del), which has been associated with X-linked Borjeson-Forssman-Lehmann syndrome (BFLS; MIM #301900); one female patient has been reported with a loss of function allele and X-inactivation (Turner et al., 2004). X-inactivation studies in our patient showed a skewed pattern, but an association with PHF6 was unlikely due to a lack of phenotypic overlap with BFLS. Furthermore, the variant identified in our patient, unlike a clear loss of function mutation reported in BFLS, leads to an in-frame deletion with no splicing defect (Fig. S9).

## Conclusions

In conclusion, we have determined novel *in vivo* functions of the OCRL adaptor proteins, PHETA1 and PHETA2. Deficiency in *pheta1/2* resulted in impaired renal physiology and craniofacial development in zebrafish, resembling the renal and craniofacial phenotypes in a UDP patient carrying a dominant-negative allele of PHETA1. The craniofacial deficits in zebrafish *pheta1/2* mutants were likely caused by a dysregulation of cathepsin K, which altered the extracellular composition of craniofacial cartilages. This results support the hypothesis that PHETA1 mutation was contributory to disease, but further studies with additional patients will be needed to determine the roles of PHETA1/2 in human disease fully.

## Materials and Methods

### Patient Enrollment, Consent, and Sample Analysis

The patient (UDP.5532) was enrolled in the National Institutes of Health (NIH) Undiagnosed Diseases Program (UDP)(Gahl et al., 2012; Gahl et al., 2015; Gahl et al., 2016) under the protocol 76- HG-0238, “Diagnosis and Treatment of Patients with Inborn Errors of Metabolism and Other Genetic Disorders”, which was approved by the Institutional Review Board of the National Human Genome Research Institute. Written informed consent was obtained from the parents of the patient.

Patient-derived fibroblasts were cultured in high glucose DMEM supplemented with Fetal Bovine Serum (15%), non essential amino acid solution, and penicillin-streptomycin with L-glutamine. Normal adult human gender matched dermal fibroblasts (ATCC PCS-201-012) were used as controls. RNA was isolated using RNeasy Mini Kit (Qiagen), and first strand cDNA was synthesized by high capacity RNA to cDNA kit (Thermo Fisher) according to the manufacturer’s protocol. For quantitative real-time PCR, primer pair specific to the three common isoforms (NM_001177996.1, NM_001177997.1, and NM_144671.4) of human *PHETA1* (Forward primer: 5’- GAAGAGCGAGCTGAGGCTG-3’, Reverse primer: 5’- GTCACAGGTGGCGTAGAAGG-3’) and housekeeping gene *POLR2A* (Forward primer: 5’-CATGTGCAGGAAACATGACA-3’, Reverse primer: 5’-GCAGAAGAAGCAGACACAGC-3’) were PCR amplified and monitored using a CFX96 Touch Real-Time PCR detection system (Bio-Rad). Relative expression of *PHETA1* transcripts was normalized to the expression of *POLR2A* and analyzed using standard delta delta Ct method. For splice site analysis of the variant in *PHF6* (NM_001015877.1:c.732_734del; p.Leu244del), we amplified the patient cDNA using *PHF6*-specific primers flanking the site of mutation and subcloned into a plasmid vector using TOPO-TA cloning (Thermo Fisher) and sequenced according to manufacturer’s instructions. Recombinant colonies were picked up by blue-white screening and extracted plasmids were sequenced using vector-specific M13 primers.

### Zebrafish Husbandry

Zebrafish of all ages were maintained under standard protocol in accordance with Institutional Animal Care and Use Committee guidelines at Augusta University, Virginia Tech, and Greenwood Genetic Center. Embryos and larvae were raised in water containing 0.1% Methylene Blue hydrate (Sigma-Aldrich). To prevent pigment formation for selected experiments, embryos were transferred to embryo media containing 0.003% 1-phenyl-2-thiourea (PTU; Sigma-Aldrich) between 18-24 hours post fertilization (hpf).

### Mutant and Transgenic Zebrafish Lines

*pheta1* (*si:ch211-193c2.2*, ZFIN ID: ZDB-GENE-041210-163, Chromosome 5: 9,677,305-9,678,075) was identified by a BLAT search using human *PHETA1* coding sequence against the UCSC zebrafish genome database (Kent, 2002). *pheta2* (*zgc:153733*, ZFIN ID: ZDB-GENE-060825-273, Chromosome 3: 32,821,205-32,831,971) was identified as a paralog of *pheta1* in the Ensembl database (Zerbino et al., 2017). Neighboring genes of *pheta1/2* was identified using the UCSC genome browser (Kent et al., 2002). Phylogenetic tree was generated using MEGA X (Kumar et al., 2018). Mutants were generated in the TL/AB mixed background using CRISPR engineering, as previously described (Jao et al., 2013; Auer et al., 2014; Gagnon et al., 2014; Irion et al., 2014).

The *pheta1*^vt2^ allele harbored a 38 base pair deletion (frame shift), resulting in the deletion of a MwoI restriction site, which was used to distinguish between wild-type and *pheta1*^vt2^ alleles. Genomic DNA flanking the deletion was amplified by PCR, followed by MwoI digestion for 2 hours at 60°C (primer sequences: cctcaaacaaactagcggacgtgtcgagta and cgcgacagagcctttacccatgattccata). After MwoI digestion, the cut wild-type bands were 230 and 300 base pairs in length, whereas the mutant band was 531 base pairs (uncut). The *pheta2^vt3^* allele harbored an 11 base pair deletion, resulting in the deletion of a NlaIII restriction site, which was used to distinguish between wild-type and *pheta2*^vt3^ alleles. Genomic DNA flanking the deletion was amplified by PCR, followed by NlaIII digestion for 2 hours at 37°C (primer sequences: ggacggtcagttctgtttctct and catgtaaacataccttcgtatcgtc). After NlaIII digestion, the cut wild-type bands were 180 and 44 base pairs in length, whereas the mutant band was 213 base pairs (uncut).

*Tg(ubi:pheta1^WT^)* and *Tg(ubi:pheta1^R6C^)* zebrafish transgenic lines were generated utilizing the Tol2-transgenesis system (Kawakami, 2007). Coding sequence for EGFP was ligated in frame to the 3’ end of the coding sequence of either the wild-type (Pheta1^WT^) or the patient-specific (*pheta1*^R6C^) Pheta1 protein, and placed into the Tol2 vector, preceded by the zebrafish *ubiquitin* promoter from the (Mosimann et al., 2011). The *Tol2-ubi:pheta1^WT^* and *Tol2-ubi:pheta1*^R6C^ vectors were then injected with Tol2 transposase mRNA into wild-type zebrafish larvae at the 1-cell stage. Potential founders were crossed to wild-type fish at 2-3 months of age and offspring was screened for EGFP-positive F1 founders.

### Whole Mount In Situ Hybridization

In situ hybridization was performed using standard protocols described previously (Prober et al., 2008; Pan et al., 2012). Sense and antisense probes were transcribed from linearized plasmid DNA using the MEGAshortscript T7 (Ambion) and mMessage mMachine SP6 (Ambion) transcription kits, respectively.

### Histochemistry and Immunohistochemistry

Alcian blue and Alizarin red staining was performed as previously described (Javidan and Schilling, 2004; Walker and Kimmel, 2007). Animals were imaged using a Nikon SMZ18 fluorescent stereomicroscope with an image capture system, and craniofacial measurements were obtained in ImageJ. Ordinary one-way ANOVA used for statistical analyses. Fluorescent images were acquired using a Nikon A1 laser scanning confocal system with a CF175 Apochromat LWD 25x water-immersion objective. Primary antibodies are as follows with dilutions: Anti-acetylated α-tubulin (Sigma, 1:1000), anti-γ-tubulin (Sigma, 1:100), znp-1 (anti-synaptotagmin2; DHSB, 1:25), zpr1 (ZIRC, 1:100), zpr3 (ZIRC, 1:100), anti-col2a1a (DHSB, 1:100), anti-GFP (Abcam, 1:1000). Alexa fluor conjugated secondary antibodies, DAPI (LifeTech), and toto-3 (Life Technologies) were used after primary antibody incubation.

### Injection of Endocytic Tracers and Analysis

Lysine-fixable 10 kDa dextran (Alexa 488 conjugated) or 500 kDa dextran (FITC conjugated) (Thermo Fisher) were prepared in PBS at 2 μg/μl final concentration. In addition, recombinant Cy3-conjugated His-tagged RAP (39 kDa), prepared in PBS at 5 μg/μl final concentration, was kindly provided by Dr. Martin Lowe (University of Manchester). Zebrafish embryos were anesthetized in tricaine (0.013% w/v; Fisher Scientific) diluted in embryo water at 72 hpf. Approximately 0.5- 1 nl of dextran or RAP was injected into the common cardinal vein using a glass micropipette and a pneumatic pressure injector (PLI90; Harvard Apparatus) and micromanipulator. Uptake in the renal tubular cells of the proximal pronephros was analyzed 1-2.5 hours post-injection (hpi), using a Nikon SMZ18 fluorescent stereomicroscope with an image capture system. Animals injected with 500 kDa Dextran were analyzed 24 hpi. Statistical analyses performed using Pearson’s chi-squared test.

### Morpholino Inhibition of OCRL Gene Expression

Morpholino (MO) knockdown of *ocrl1* was performed as previously described (Coon et al., 2012), and was kindly provided by Dr. Martin Lowe (University of Manchester). A translation blocking MO was utilized and 1-5 ng/ul was injected into embryos at the 1-cell stage (sequence: AATCCCAAATGAAGGTTCCATCATG). The specificity of this MO has previously been validated by rescue *ocrl1* mRNA experiments (Coon et al., 2012).

### Cilia and Craniofacial Quantification/Analysis

Cilia in the anterior portion (just anterior to the yolk extension) and posterior portion (near the cloacae) of the pronephros in the zebrafish larvae were selected and analyzed. The number of cilia within a 100×100 micron area were quantified, and the length of five randomly selected cilia were measured within the area. Craniofacial morphometric measurements were obtained with Fiji (Schindelin et al., 2012). Type II collagen was quantified by mean fluorescence intensity within a 2500 μm^2^ area in the ceratohyal cartilage and a 1000 μm^2^ area in Meckel’s cartilage.

### BMV109 Delivery and in-gel analyses

The BMV109 fluorescent ABP was injected into 3 dpf larvae (1 nl at 10 µM) pericardially via microinjection. This equates to a final global concentration of 10 nM. Probe was circulated over night at 28.8°C and harvested at 15 hpi. 25 larvae per condition were collected and lysed in citrate buffer (50 mM citrate buffer pH5.5, 5 mM DTT, 0.5% CHAPS, 0.75% Triton X-100) by brief sonication. Samples were centrifuged for 15 minutes at 15,000g and the supernatant collected. Protein concentration was determined via a micro BCA assay (cat#23235; Thermo Fisher) and samples run on 4-20% precast gradient gels containing the “stain free” tri-halo compound (Bio-Rad). UV light activated tri-halo covalently binds tryptophan residues. Equivalent protein loads were evaluated on a Bio-Rad Chemidoc MP Imaging System using this stain-free method. BMV109 Cy5 fluorescence was subsequently analyzed in gel. Total protein load per lane and individual ABP- reactive bands were quantitated using Chemidoc MP software. Individual ABP-reactive bands were normalized to total protein load and the fold difference calculated between WT and MLII samples. Gel images were processed on Adobe Photoshop (CS6 extended, version 13.0).

### Pharmacological inhibition

Cathepsin K activity was inhibited from 3 to 4 dpf in live embryos by introducing the 50 nM odanacatib (solubilized in DMSO) directly into their growth media. In all cases, WT control larvae were treated with an equivalent amount of DMSO (0.1%).

### Cell Culture

HeLa cells (ATCC CCL-2) were grown in DMEM supplemented with 5% FBS (Thermo Fisher) and 1% penicillin-streptomycin (Life Technologies). Cells were transfected using Effectene (Qiagen) according to instructions provided by the manufacturer. The FAM109A cDNA was synthesized by Genewiz and cloned into the pEGFP-C3 vector (Promega). The R6C mutation was introduced into the construct using the Q5 site-directed mutagenesis kit (NEB). The pcDNA3-HA-human OCRL plasmid was a gift from Pietro De Camilli (Addgene plasmid # 22207; http://n2t.net/addgene:22207; RRID:Addgene_22207).

### Protein-protein Interaction

Lysates were prepared from transfected HeLa cells by incubating the cell pellet in RIPA buffer (50 mM Tris-Cl pH 7.5, 150 nM NaCl, 1% NP40, 1 mM EDTA). The lysate was clarified by centrifugation at 10,000g for 5 minutes at 40C. 1x Halt protease inhibitor cocktail (Pierce) was added to the lysate. 800 μg total protein was used per immunoprecipitation. Immunoprecipitation was done using GFP-Trap beads (Chromotek) in binding buffer (50 mM Tris pH 7.5, 150 mM NaCl, 0.2 mM EDTA, 0.05% NP40). The lysate was incubated with beads for 75 min at 40°C. Subsequently, the beads were washes four times using binding buffer. The bound proteins were eluted by boiling in Laemmli buffer, run on a gel and analyzed by western blotting. A monoclonal GFP antibody (JL-8; Clontech) and a monoclonal HA antibody (sc-7392; Santa Cruz) were used in the western. The HRP signal was acquired on a Chemidoc MP (Bio-Rad) imaging system.

### qRT-PCR Analysis of Transcript Abundance in Zebrafish

Total RNA was isolated from four pools of five larvae (head only) using the RNA Miniprep Kit (Zymo). First-strand cDNA synthesis was performed using the SuperScript III First-Strand Synthesis System (Thermo) with 50 ng of total RNA. A threefold dilution of the cDNA reaction was used as template for the qRT-PCR reactions. Primer sequences were used as previously described (Petrey et al., 2012). qRT-PCR reactions were run in technical duplicates, and reactions consisted of 1 μl diluted cDNA, 2 μl of gene-specific primer pair and 5 μl iTaq Universal SYBR Green Supermix (Bio-Rad). The relative transcript abundance of each gene (normalized to *rpl4*, a housekeeping gene) for each group of pooled samples were determined and analyzed.

### Optokinetic Response in Zebrafish

VisioTracker 302060 (New Behavior TSE) was used for OKR assay. Eye movements of individual fish were recorded at 5 frames per second by an overhead CCD camera. Zebrafish larvae were placed in the center of an uncoated 50mm glass bottom petri dish (MatTek) and immobilized in 1.5-2% low melting agarose (Fisher Scientific) in E3 buffer. Agarose around the eye was removed with forceps to allow free eye movement. The dish was then filled with water. To test slow phase performance under short periodicity, direction of black and white grating switched every three seconds with grating velocity at 7.5°/s. Each experimental run (trial) was 108 seconds long and included twelve 9-second phases at varying contrast levels (0.99, 1.0, 0.5, 0.2, 0.1, 0.05, 0.02, 0.05, 0.1, 0.2, 0.5, 1.0). 5-6 trials were tested for each animal. Contrast sensitivity and eye correlation were calculated in trials where behavioral response is robust.

### Image Processing and Statistical Analyses

Images were processed with Fiji (Schindelin et al., 2012) and Photoshop (Adobe Systems) software. All statistical analyses were performed in GraphPad Prism (Version 7.0d). The chi-square test, one-way, and two-way ANOVA test was performed as appropriate. All values are expressed as mean ± SEM, unless otherwise noted. The test was considered significant when p<0.05. When ANOVA tests were found to be significant, the Holm Sidak’s post-hoc test was performed to make pairwise comparisons.

## Acknowledgements

This work was supported by funding from the National Institutes of Health (GM119016 to KMA, HK, GBG, and YAP; GM086524 to TM, TW, and HFS), the Medical College of Georgia, and Virginia Tech. We thank the animal care staff at Augusta University and Virginia Tech for animal husbandry, the clinicians and scientists at National Human Genome Research Institute for the clinical data, members of the Pan Laboratory, Rachel Roberts and Michelle Ma, for training in histological and behavioral assays, and M. Lowe (University of Manchester, London) for his assistance with the endocytic assays. Lastly, we are grateful to the patient and her family for their participation in the Undiagnosed Diseases Program.

## Author Contributions

KMA and YAP conceived the study, with input from GBG and HFS. KMA, TW, and YAP generated the *pheta1^vt2^* and *pheta2^vt3^* mutant lines, as well as the *pheta1^R6C^* transgenic lines. KMA, TW, HFS, and TM contributed to the histological, morphological, and endocytosis analyses. RVK and GBG performed the coimmunoprecipitation assays. KMA performed the behavioral analyses. LAW, DA, TM, CJT, JS, MCM, and WAG contributed to patient variant identification and clinical evaluations of the UDP patient. HGK, WW, and PA contributed to the protein modeling of the R6C mutation in PHETA1. KMA and YAP wrote the manuscript, with contributions from HFS.

---

**Supplementary Figure S1.**
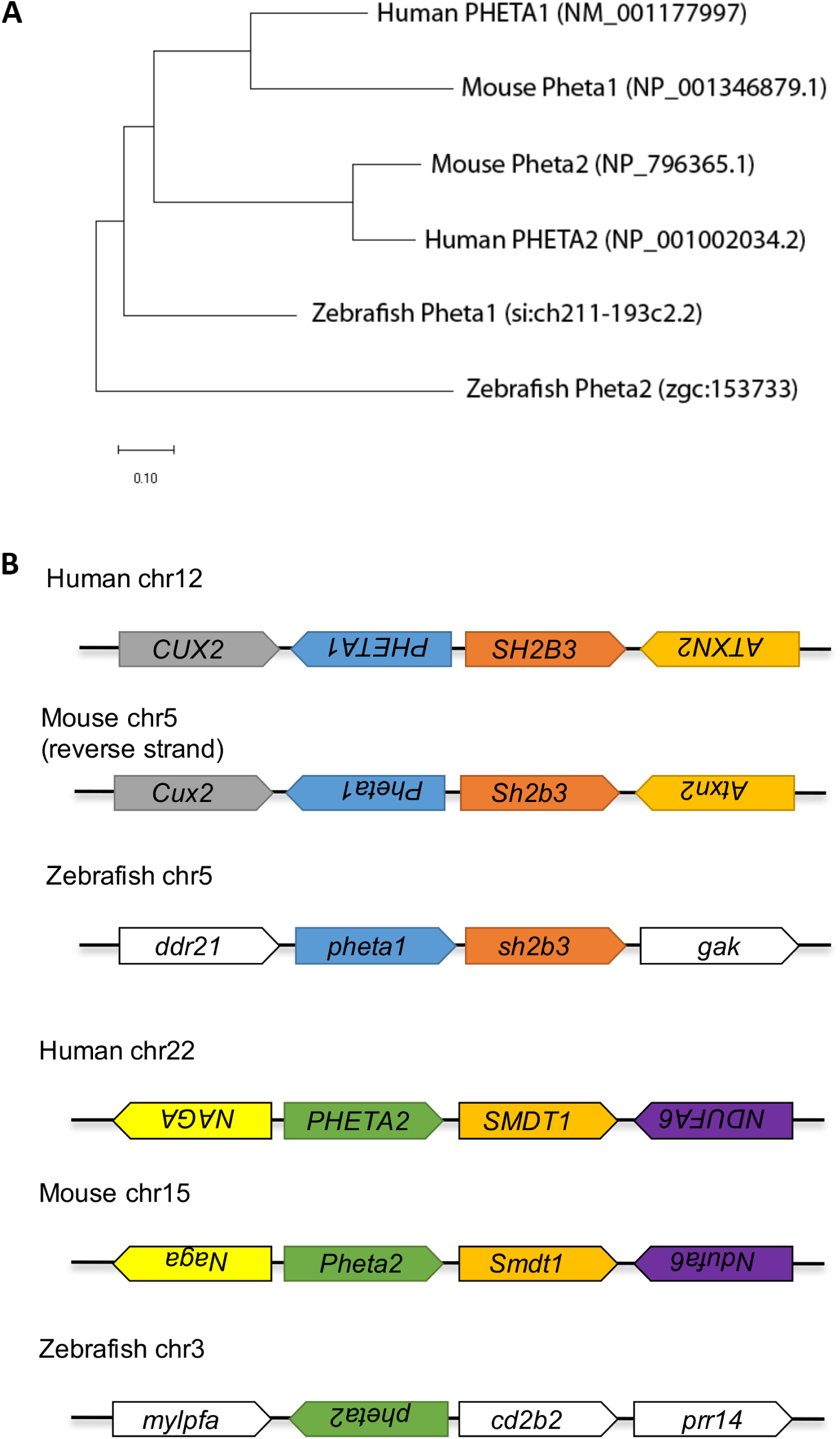
Evolutionary relationship of human *PHETA1*/2 with mouse and zebrafish homologs. (A) The evolutionary history was inferred using the Neighbor-Joining method. The evolutionary distances were computed using the Poisson correction method and are in the units of the number of amino acid substitutions per site. Evolutionary analyses were conducted in MEGA X. (B) Genomic organization of human *PHETA1/2* and their respective homologs in mouse and zebrafish.

**Supplementary Figure S2.**
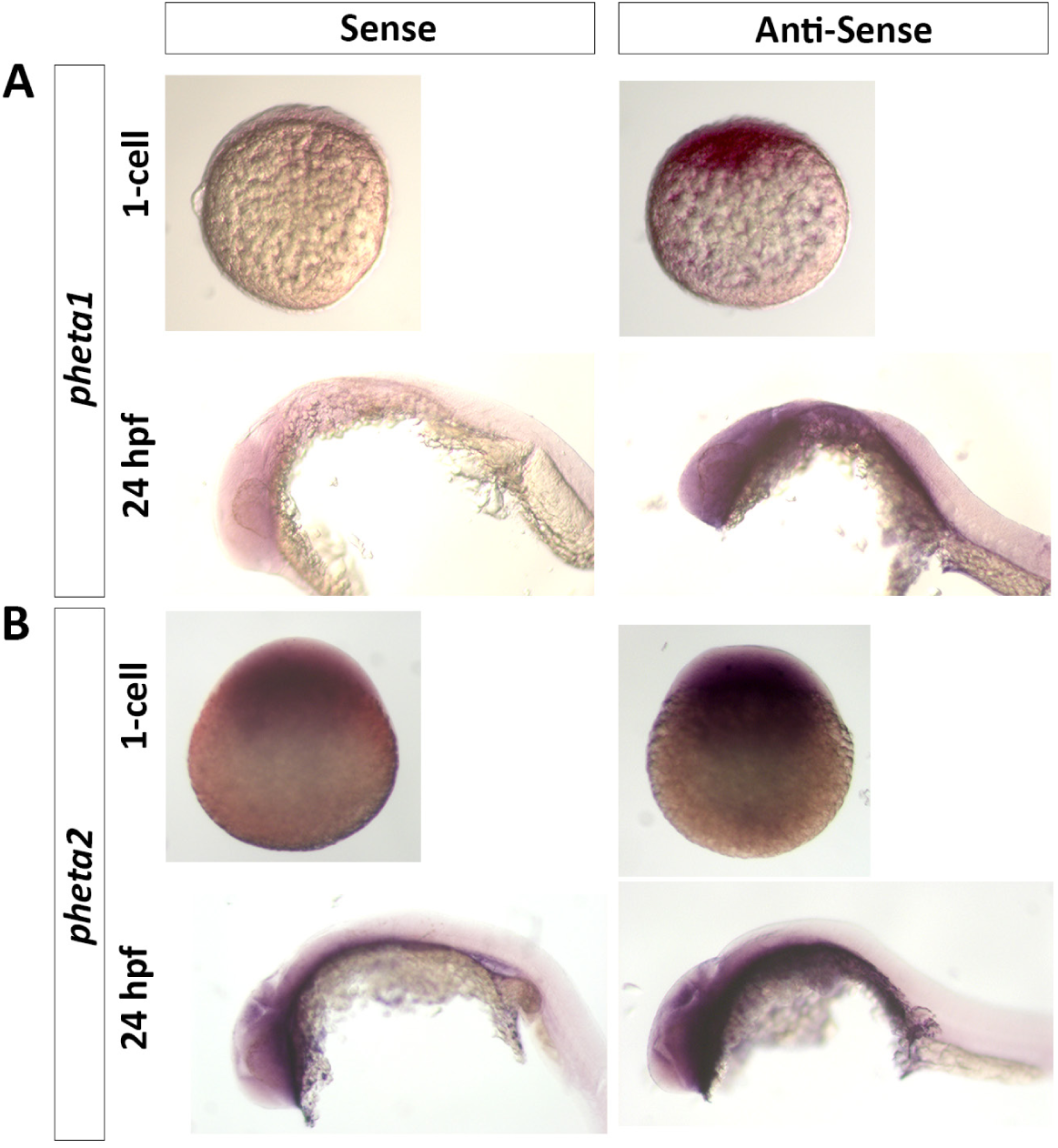
*pheta1* and *pheta2* whole-mount *in situ* hybridization of developing zebrafish embryo. Wild-type embryos were collected and whole-mount *in situ* hybridization performed at the 1-cell stage and 24 hours post-fertilization (hpf). Hybridization with anti-sense probe showed that *pheta1* (A) and *pheta2* (B) are expressed maternally (1-cell) and during development (24 hpf). Sense probe labeling serves as the negative control.

**Supplementary Figure S3.**
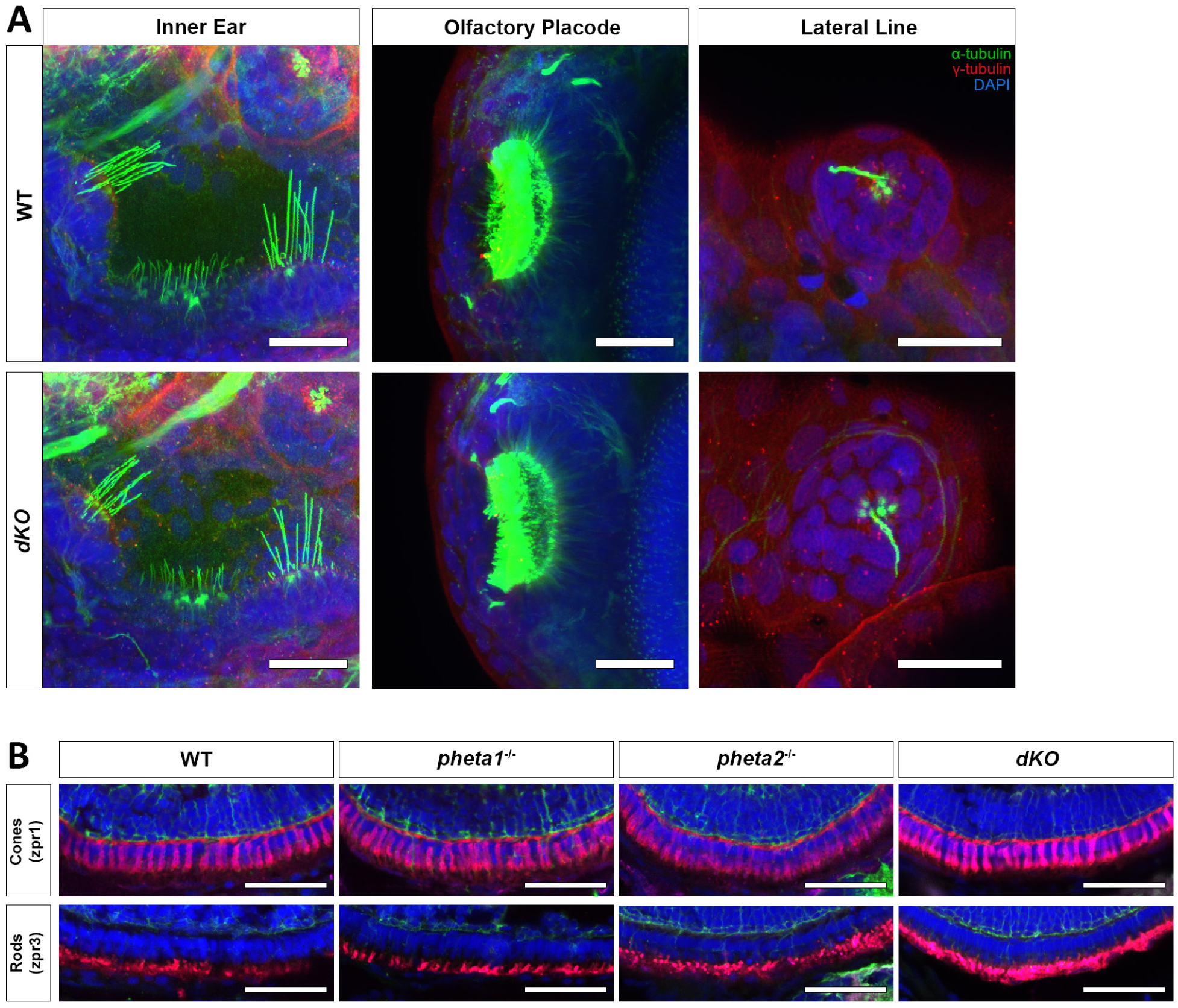
Loss of *pheta1/2* has no effect on ciliogenesis in the inner ear, olfactory placode, lateral line, and photoreceptors. (A) Representative confocal images of cilia in WT and *dKO* in three organs are shown: the inner ear, the olfactory placode, the lateral line, and the anterior pronephros. Cilia are labeled with anti-acetylated α-tubulin (green), basal bodies labeled with anti-γ tubulin (red), and nuclei labeled with DAPI (blue). Scale bars= 25 µm. (B) Confocal images of transverse cryosections obtained from 5 dpf larvae. Antibodies include DAPI (blue) and acetylated α-tubulin (green). Zpr1 (red) and zpr3 (red) were used to label cones and rods, respectively. Scale bars= 50 µm.

**Supplementary Figure S4.**
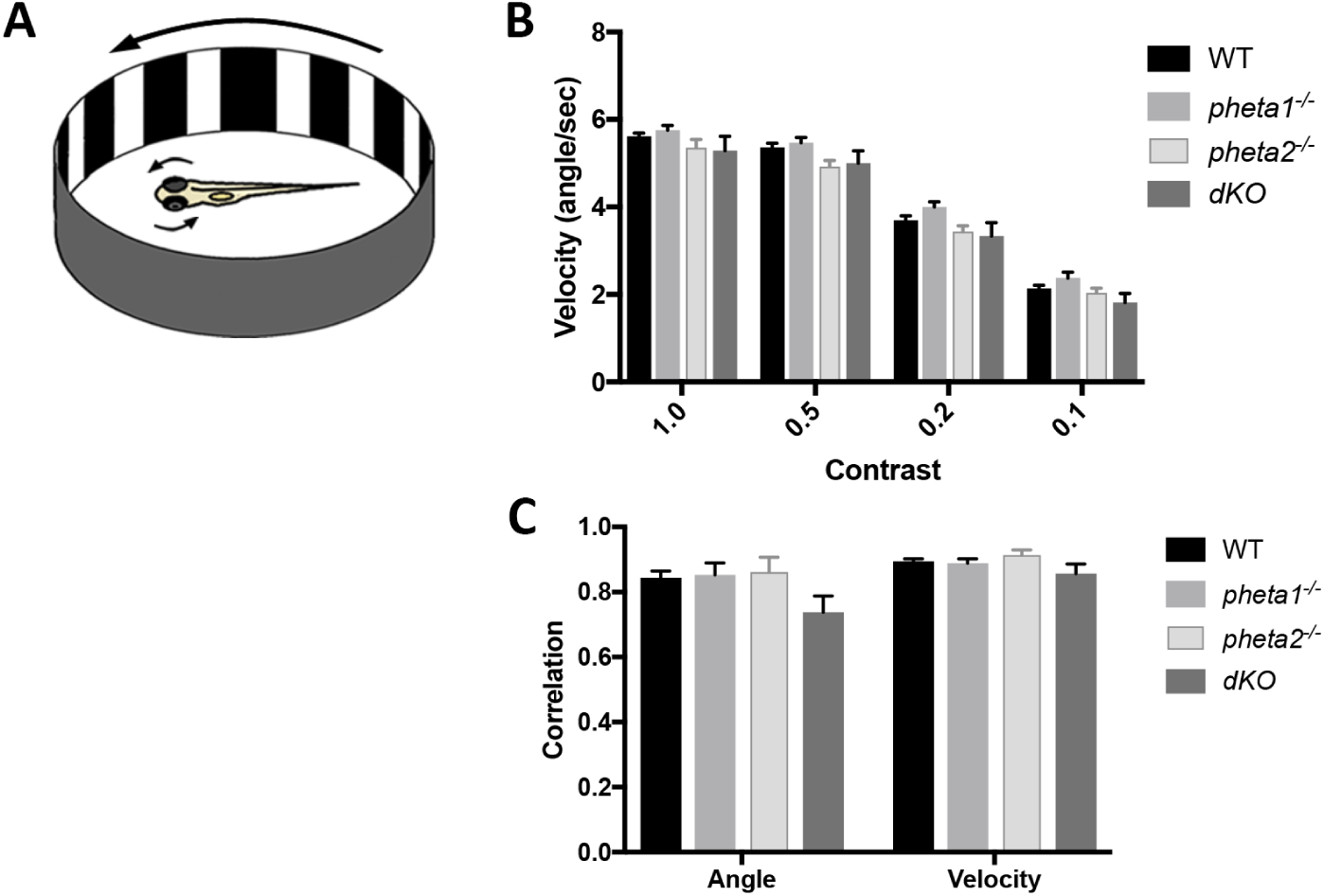
Loss of *pheta1/2* has no effect on OKR. (A) A zebrafish larva is placed in an arena with moving black and white gratings. An infrared camera records the position and speed of each eye during the visual stimulation. (B) Velocity of tracking movements in response to moving gradients at various contrasts; 1.0, 0.5, 0.2, 0.1. Error=SEM. (C) Correlation in angle and velocity between left and right eye. WT: n=34, *pheta1*^-/-^: n=18, *pheta2*^-/-^: n=8, *dKO*: n=10. Statistical analyses performed using two-way ANOVA with Holm Sidak post-test.

**Supplementary Figure S5.**
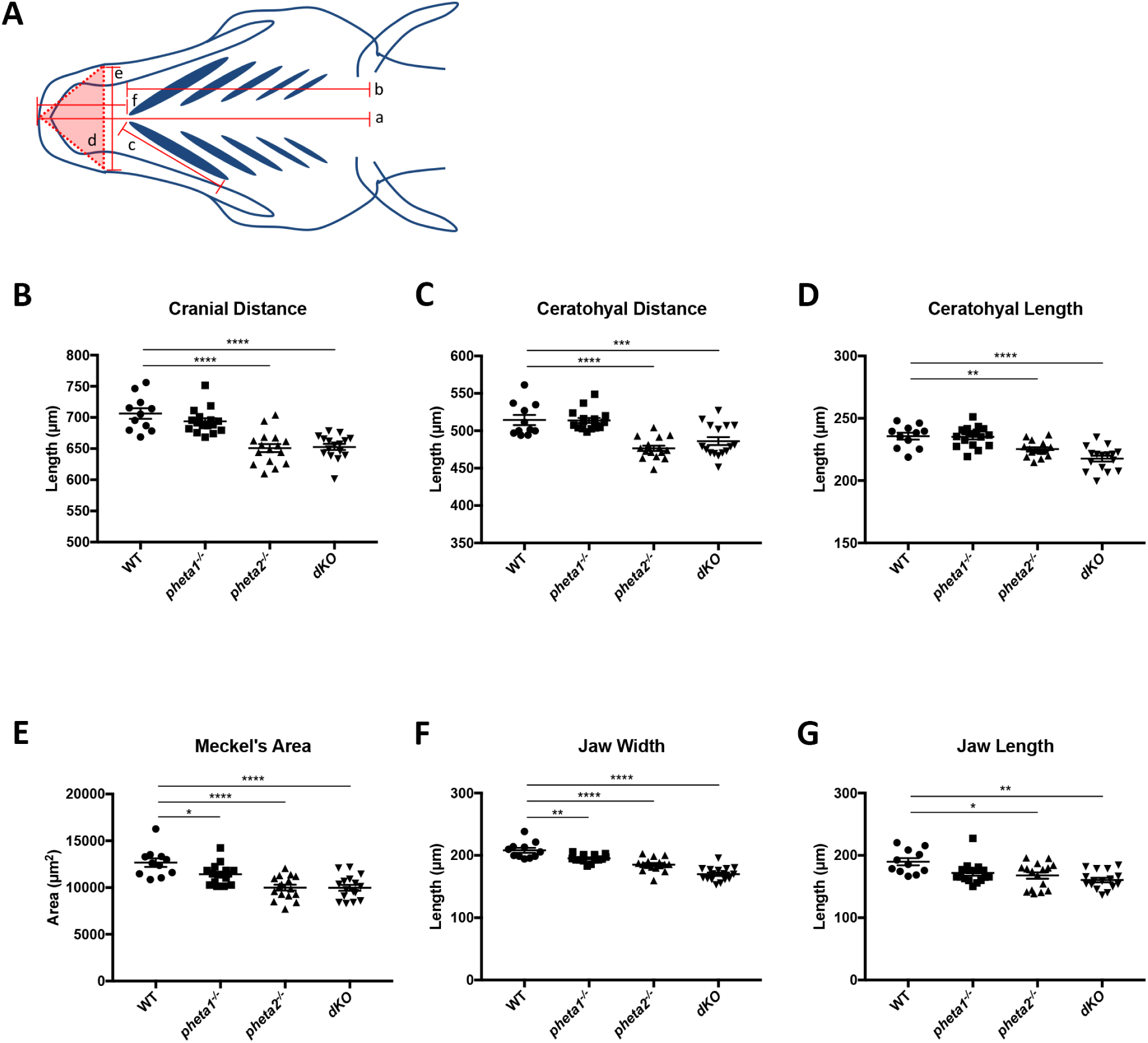
Morphometric measurements of craniofacial development in WT, *pheta1^-/-^*, *pheta2^-/-^*, and *dKO*. (A) Schematic with measurements analyzed. a: cranial distance (B); b: ceratohyal distance (C); c: ceratohyal length (D); d: Meckel’s area (E); e: jaw width (F); f: jaw length (G). Error=SEM. Statistical analyses performed using one-way ANOVA with Holm Sidak post-test. WT: n=11, *pheta1^-/-^*: n=16, *pheta2^-/-^*: n=16, *dKO*: n=16. *p<0.05, **p<0.01, ***p<0.001, ****p<0.0001.

**Supplementary Figure S6.**
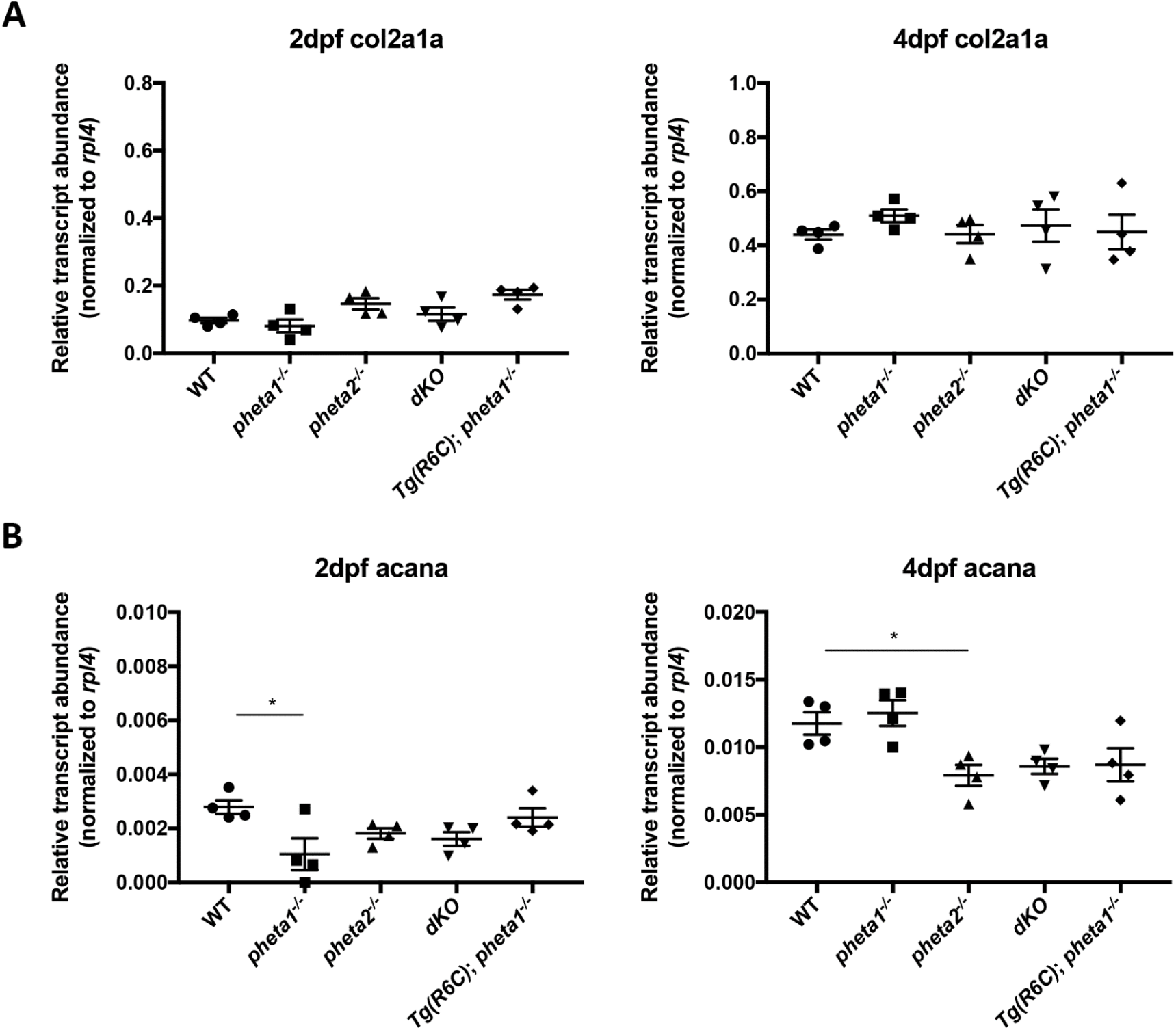
qRT-PCR of *col2a1a* and *acana* in 2dpf and 4dpf zebrafish larvae. (A) Relative transcript abundance of *col2a1a*. (B) Relative transcript abundance of *acana*. Statistical analyses performed using one-way ANOVA with Holm Sidak post-test. n=4 groups of pooled samples per genotype. *p<0.05.

**Supplementary Figure S7.**
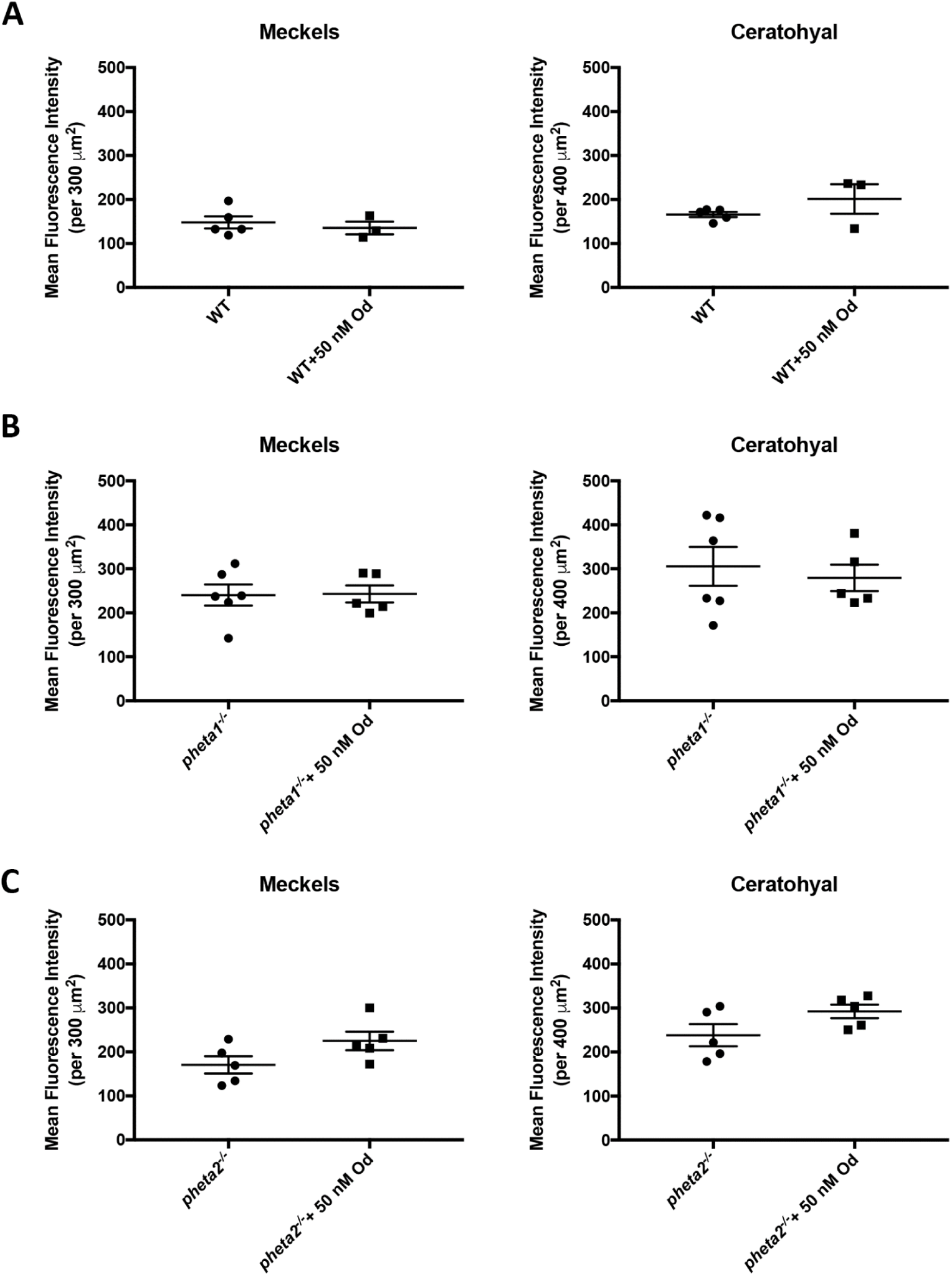
Type II collagen immunohistochemistry after administration of odanacatib. Quantification of mean fluorescence intensity in the Meckel’s and Ceratohyal cartilage for (A) WT animals, (B) *pheta1****^-/-^***animals, and (C) *pheta2****^-/-^***animals. Od: odanacatib. Statistical analyses performed using one-way ANOVA. n=5-6 animals per genotype. *p<0.05, **p<0.01.

**Supplementary Figure S8.**
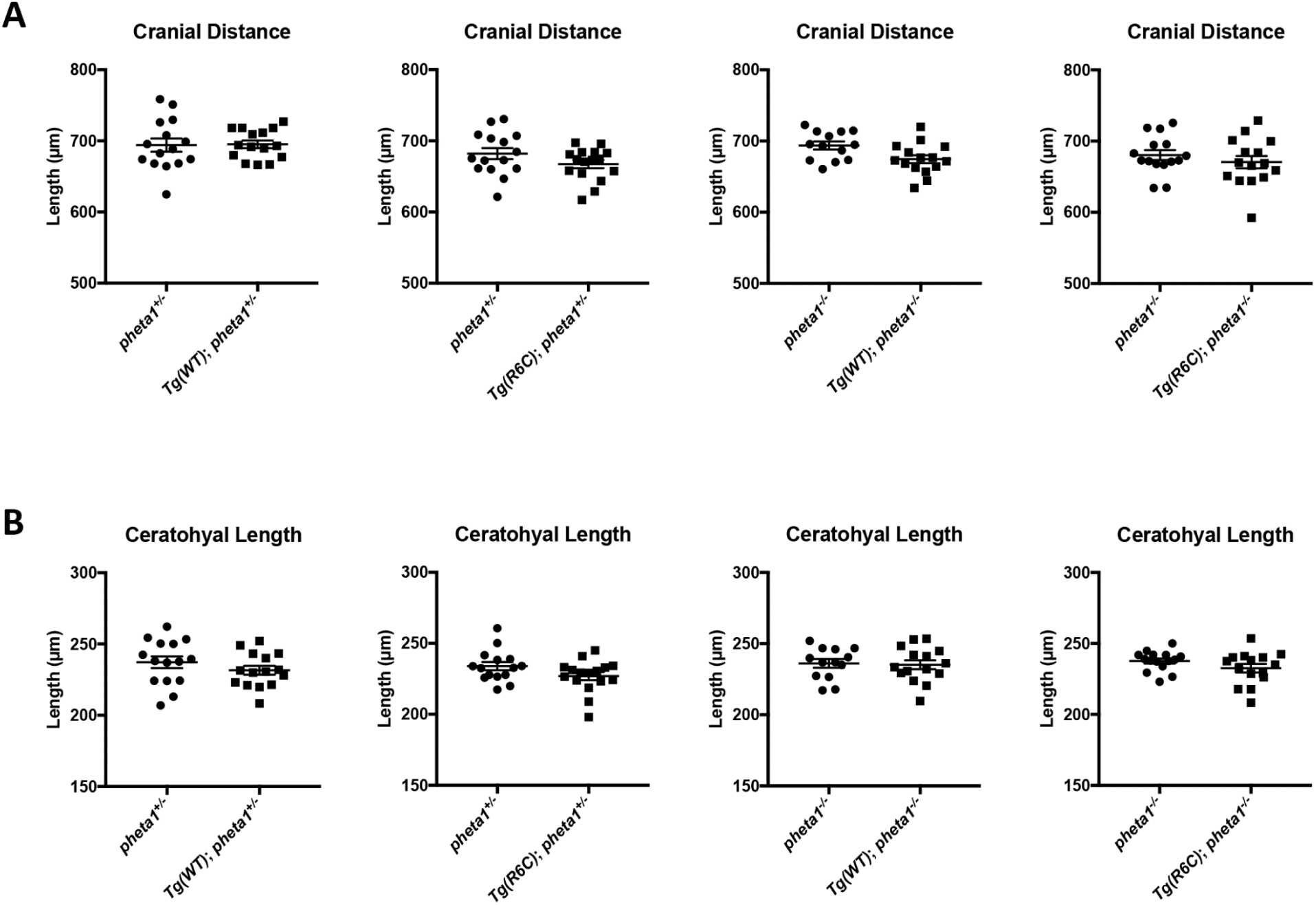
Effect of *Tg(WT)* and *Tg(R6C)* on cranial distance and ceratohyal length. (A) Cranial distance in *pheta1*^+/-^ or *pheta1*^-/-^ background. (B) Ceratohyal length in *pheta1*^+/-^ background or *pheta1* ^-/-^ background. Statistical analyses performed using one-way ANOVA with Holm Sidak’s multiple comparisons. n=15-16 animals per condition.

**Supplementary Figure S9.**
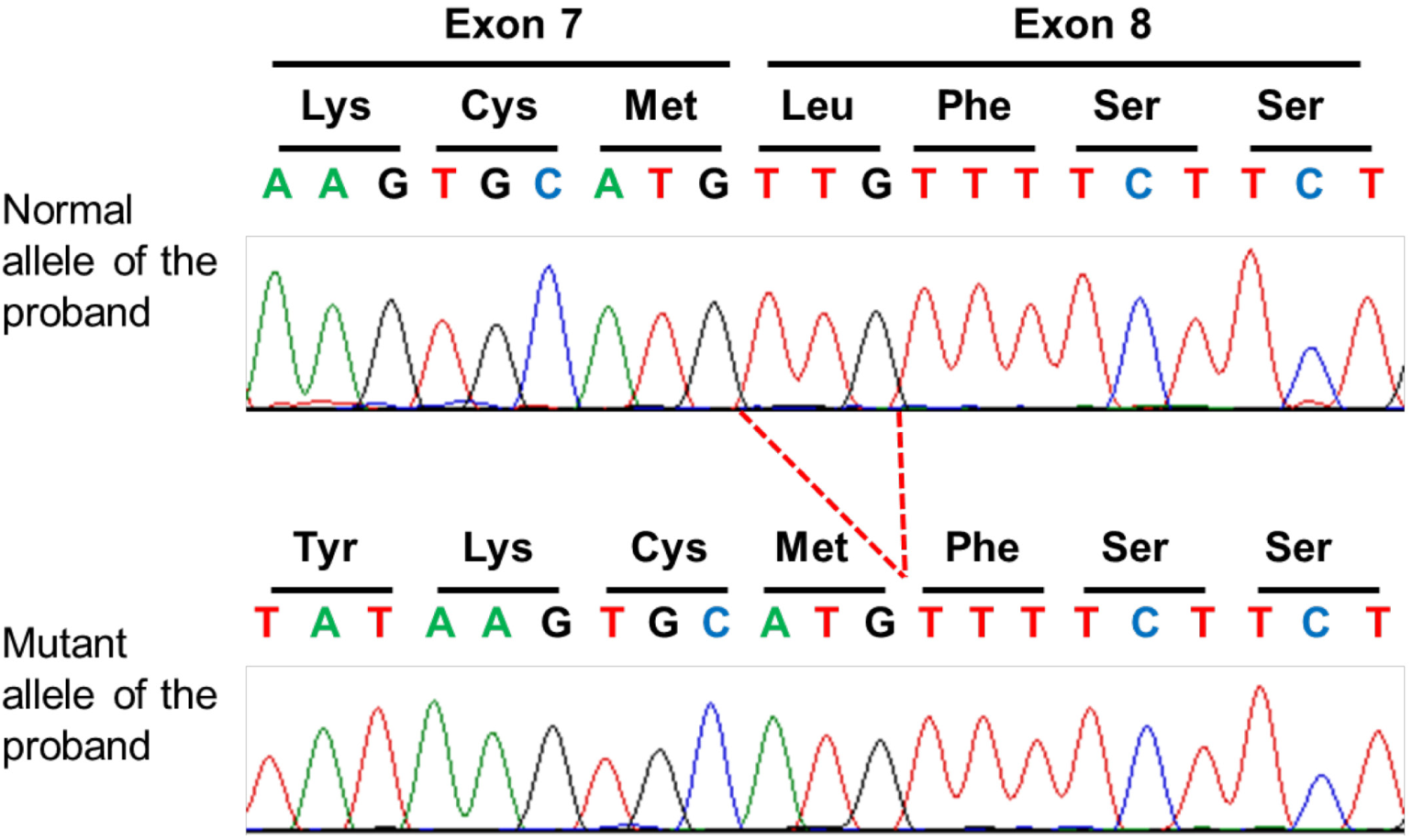
Splice site analysis of the *PHF6* variant in the UDP patient. Sequence chromatograms showing the normal allele (upper panel) and mutant allele (lower panel) of the UDP patient (i.e., proband). There was no splice defect except the in frame deletion of Leu244 at the exon 7 and 8 boundary (marked in red).

